# AI-Driven Virtual Screening and First-in-Class Conformational Biosensor Enable the Discovery of a GPR183 Inverse Agonist

**DOI:** 10.1101/2025.06.19.659764

**Authors:** Louise Andersson, Michele Roggia, Kittikorn Wangriatisak, Maria Gil, Karine Chemin, Sandro Cosconati, Paweł Kozielewicz

**Affiliations:** Molecular Pharmacology of GPCRs, Department of Physiology and Pharmacology, Karolinska Institutet, 171 65 Solna, Sweden; DiSTABiF, University of Campania Luigi Vanvitelli, Via Vivaldi, 43, 81100 Caserta, Italy; Division of Rheumatology, Department of Medicine, Solna, Karolinska Institutet, Karolinska University Hospital, Stockholm, Sweden; Center for Molecular Medicine, Karolinska Institutet, 171 76 Solna, Stockholm, Sweden

**Author notes:** equal contribution (Louise Andersson, Michele Roggia). Corresponding Authors Paweł Kozielewicz - Molecular Pharmacology of GPCRs, Department of Physiology and Pharmacology, Karolinska Institutet, 171 65 Solna, Sweden Sandro Cosconati - DiSTABiF, University of Campania Luigi Vanvitelli, Via Vivaldi, 43, 81100 Caserta, Italy.

**Keywords:** G protein-coupled receptors, GPR183, Artificial Intelligence-driven ligand screen, conformational sensors, inverse agonist, migrationw

## Abstract

GPR183 is a chemotactic G protein-coupled receptor implicated in immune cell migration and various diseases. Here, we report the discovery of novel GPR183 inverse agonists using an artificial intelligence-driven virtual screening approach combined with biophysical sensors including a first-in-class conformational biosensor. From an initial screen of 70 compounds and a hit expansion, a selected candidate - compound 78 - demonstrated potent inhibition of GPR183’s constitutive and agonist-stimulated Gi activation as well β-arrestin2 recruitment. The binding of the compound to the receptor core was confirmed by studies using a new conformational sensor based on intramolecular BRET, molecular dynamics, and site-directed mutagenesis experiments. Finally, the compound 78 was able to block agonist-induced migration of peripheral blood mononuclear cells *ex vivo*. This work provides a novel chemical scaffold for GPR183 modulation. It introduces a new tool for dissecting receptor function, paving the way for therapeutic exploration in inflammatory, autoimmune, and neoplastic disorders.

## INTRODUCTION

G protein-coupled receptors (GPCRs) constitute the largest family of membrane proteins in the human genome, represent the key mediators of signalling responses, and are the largest group of targets for marketed drugs [1]. Among them, GPR183 (also known as Epstein-Barr virus-induced gene 2, EBI2) is primarily a Gi-coupled receptor [2], initially identified through its upregulation in B cells infected with Epstein-Barr virus [3], involved in the guidance of B cells, T cells and dendritic cells, within lymphoid and non-lymphoid tissues [4–6]. To this end, GPR183 is recognised as a chemotactic receptor for oxysterols [7, 8], with 7α,25-dihydroxycholesterol having the highest efficacy, potency, and binding affinity at the receptor [2, 8]. While the efficacy of the endogenous ligand has been validated for nanomolar concentrations for this agonist in multiple studies from different laboratories [2, 9–13], at concentration levels corresponding to physiological of oxysterols [14], the receptor is still classified as orphan by the International Union of Basic and Clinical Pharmacology (IUPHAR). In addition to the ligand-induced signalling, constitutive activity of GPR183 in the apparent absence of ligand stimulation has been documented following receptor plasmid DNA overexpression as well as at endogenous levels of the receptor’s expression [15].

Mounting evidence continues to implicate GPR183 in various physiological and pathological processes. To this end, high expression and/or activation of GPR183 has been proposed to play a role in a range of diseases, including cancer [16], metabolic disorders [17], inflammatory [18], and autoimmune diseases [19]. Importantly, GPR183 expression and activity appear to play context-dependent roles - beneficial in some settings, such as diffuse large B-cell lymphoma [16], and detrimental in others, such as in acute myeloid leukaemia [20]. As such, pharmacological modulation of GPR183 is of growing interest, with both synthetic agonists [10, 21] and inverse agonists [22–26] being explored as potential therapeutic tools. Please note that, taking into consideration the dynamic nature of GPCRs and of GPCR-ligand interaction [27], an observation that pure antagonists are in fact rare [28] and already-mentioned constitutive activity of GPR183, we assume that a lot of the supposed GPR183 antagonists will exhibit negative efficacy (more or less pronounced depending on an assay) and as such, we will refer to these compounds simply as inverse agonists.

The pursuit of effective GPR183 modulators for treating inflammatory and autoimmune diseases has led to the development of numerous distinct chemical classes of inverse agonists. Early efforts established the piperazine diamide scaffold, exemplified by NIBR189 [22], a selective and orally bioavailable inverse agonist with an *IC*_50_ of approximately 11 nM that demonstrated preclinical efficacy in models of inflammatory bowel disease (IBD) and viral infections. Another key compound from this class is ML401 [29], a potent inverse agonist (*IC*₅₀ ≈ 1 nM) with favourable ADME properties, widely used as a research tool. However, challenges associated with this scaffold, including off-target effects such as hERG inhibition and suboptimal pharmacokinetic profiles, spurred further optimization. This led to the creation of second-generation compounds like the recently reported a benzo[d]thiazole derivative [24], which achieved sub-nanomolar potency (*IC*₅₀ = 0.82 nM) while significantly improving the hERG safety window and *in vivo* efficacy in colitis models.

In parallel, structurally distinct chemotypes have emerged, indicating a broadening of the chemical space for GPR183 modulation. The spirocyclic inverse agonist GSK682753A (*IC*₅₀ ≈ 54 nM) [30] was instrumental in elucidating the receptor’s inactive state through cryo-EM studies. Despite its utility as a tool compound, its poor metabolic stability has limited its therapeutic potential. More recently, a novel class of difluoro-1,3-benzodioxole-piperazine-pyrimidine inverse agonists was disclosed, with a lead compound (HY-162011) showing excellent potency (β-arrestin *IC*₅₀ = 8.6 nM) [23], a clean safety profile, and high oral bioavailability, proving effective in a preclinical model of rheumatoid arthritis. While these represent significant advances, the existing scaffolds still present limitations, whether in specific ADME properties, off-target liabilities, or synthetic complexity. This landscape underscores a clear and ongoing need for the discovery of new, structurally diverse GPR183 modulators with optimized, drug-like properties to fully exploit the therapeutic potential of this target.

To address the critical need for novel chemical scaffolds, we moved beyond conventional discovery pipelines and instead exploited the predictive power of advanced computational chemistry. We hypothesized that an Artificial Intelligence (AI)-powered Structure-Based Virtual Screening (SBVS) could rapidly and effectively identify unique GPR183 modulators that traditional screening campaigns might overlook. In particular, in this study we implemented a the PyRMD2Dock approach developed by some of us, which leverages the classification power of the Random Matrix Discriminant (RMD) algorithm to enhance the throughput of virtual screening campaigns for drug discovery [31]. These studies were coupled with the use of established bioluminescence resonance-energy transfer (BRET)-based biophysical sensors (biosensors) and a novel, first-in-class conformational biosensor to identify GPR183 inverse agonists. These studies led to the discovery of novel chemotypes that were further evaluated in cell-based migration assays to probe the inhibition of agonist-stimulated GPR183-mediated signalling. These findings establish a new tool for probing GPR183 function and provide a foundation for therapeutic development targeting this receptor.

## MATERIALS AND METHODS

### Molecular docking

Cryo-EM structure of GPR183 in complex with its endogenous ligand 7α,25-dihydroxycholesterol (PDB 7TUZ) [2] was sourced from the RCSB PDB database and underwent preliminary adjustments for docking purposes using the protein preparation wizard integrated into the Schrödinger suite [32]. The hydrogen atoms were added and minimized, the solvent molecules were removed, and the appropriate protonation and tautomeric state of the protein side chains were calculated at physiological pH. Using the AutoDockTools Python [33] scripts, the GPR183 structure was converted into the AutoDock PDBQT format, where, compared to a standard PDB file, Gasteiger charges are added to the atoms and the torsional freedoms of the various bonds are described. Then, the receptor grid maps were calculated with the AutoGrid4 software, mapping the receptor interaction energies using every AutoDock atom type as a probe and the docking grid box was centred on the binding site on the 7α,25-dihydroxycholesterol structure (XYZ size of the box: 60 x 60 x 60 with a spacing of 0.375 Å). From the ZINC20 database, a set of 1M randomly chosen compounds was selected and the molecules’ SMILES strings were converted into 3D structures with the employment of the LigPrep (Schrödinger Release 2024-1: LigPrep, Schrödinger, LLC, New York, NY, 2024.) routine in Schrödinger’s Maestro suite. Their tautomeric and protonation states were calculated at physiological pH, and the possible enantiomers were generated. The prepared molecules were exported as PDB files and converted into PDBQTs by making use of the AutoDock suite scripts [33]. For the docking calculations, attained through AD4-GPU [34], the Lamarckian Genetic Algorithm (LGA) was employed, encompassing a total of 50 LGA runs. All other settings were maintained at their default values. The docking results were subsequently grouped based on the RMSD criterion, whereby solutions differing by less than 2.0 Å were considered part of the same cluster. The ranking of these clusters was determined based on the calculated free energy of binding (ΔG_AD4_).

### AI-enforced VS

To create a ML prediction model, PyRMD was fed with a comma separated file (.csv) generated by extracting the predicted lowest Δ*G*_AD4_ for each of the 1M docked compounds randomly chosen from the ZINC20 medium-large database. This led to the construction and preparation of the training dataset. According to their predicted Δ*G*_AD4_, the compounds included in the .csv file were classified into three groups: “actives”, “inactives”, and discarded. Compounds who’s predicted Δ*G*_AD4_ falls below the “activity” thresholds of -9.5, -10.0, -10.5, -10.75, and -11.0 kcal/mol were placed in the actives group. Instead, those with Δ*G*_AD4_ value higher than the “inactivity” threshold of -4.0, -4.5, -5.0, -5.5, and -6.0 kcal/mol went into the “inactives” group. Moreover, compounds whose Δ*G*_AD4_ value falls above the “activity” threshold and “below-the-inactivity” threshold were discarded. By selecting the MinHash fingerprints (MHFP) for the featurization process, and by varying the ε cutoff for “actives” (0.01-0.99 with a 0.10 step) and “inactives” (0.01-0.99 with a 0.10 step), 3025 different models were generated. For all the models PyRMD returns relevant metrics to evaluate their predictive capability (i.e., TPR, FPR, F-Score, ROC AUC, BED ROC, PRC AUC). In this work, the selected model was chosen by maximizing the True positive rate (TPR)/False positive rate (FPR) trade-off. Once the model was generated, PyRMD was used to screen the remaining ∼9M compounds from ZINC20 database and it automatically returns all the compounds deemed to be active along with a confidence score of its prediction (RMD Score).

### Molecular dynamics

The ligand-receptor complex obtained from the docking experiments was used to construct a molecular dynamics system. Initially, the complex was embedded in a 1-palmitoyl-2-oleoylphosphatidylcholine (POPC) membrane and solvated in water within an orthorhombic system with a buffer distance of 10 Å using CHARMM-GUI [35]. Additionally, the overall charge of the system was neutralized, and the salt concentration was set to 0.15 M KCl. Missing Cryo-EM protein’s extracellular loops (ECL1 and ECL2) were added using Alphafold [36, 37] (Uniprot [38] code: P32249). To minimize the fixed structure and adapt the added loops to the initial protein, 2500 steps with the Steepest Descent method followed by 5000 steps employing a Polak-Ribier Conjugate Gradient were applied, adding constrains on the non-reconstructed residues. The complex underwent molecular dynamic simulation using Amber22 (Amber manual, Amber 2025, University of California, San Francisco) The general AMBER force field (gaff2) [39] was applied to **78** through antechamber [40], while the protein force field (ff19SB) [41] was employed for the GPR183 structure to create its topology parameters, using tleap [42]. To stabilize the water-counterion system, energy minimization was conducted. Initially, the solvent’s energy and positions were adjusted through 50000 steps of steepest descent energy minimization, followed by an additional 10000 steps of conjugate gradient minimization. Meanwhile, the complex and the membrane remained fixed with a constraint of 10.0 kcal·mol^-1^·Å^-2^. Second, the entire system was gradually heated from 0 K to 300 K over 3 different steps of 20 ps each with position restraints applied at constant volume. The SHAKE algorithm was utilized to constrain covalent bonds involving hydrogen atoms, with a relative geometrical tolerance of 0.00001. Subsequent other 2 NVT equilibration steps were applied at constat temperature (300K), gradually relaxing the protein and membrane constraints encompassing a total of 160ps. Then, 4 different isothermal isobaric ensemble (NPT)-MD steps were performed for a total of 850 ps to equilibrate the system at 300 K and 1 bar gradually removing the constraints to relax the system (last 500ps without constrains). Finally, a 500ns simulation was performed maintaining a constant temperature of 300 K and pressure of 1 bar. The trajectory was updated every 5000 fs for further analysis. Analysis of the MD trajectory was attained with the cpptraj library [43] within AmberTools23.

### In vitro cell culture

HEK293A cells (Thermo Fisher Scientific) were cultured in DMEM supplemented with 10% FBS (Sigma), 1% penicillin/streptomycin, 1% L-glutamine (both from Thermo Fisher Scientific) in a humidified CO_2_ incubator at 37°C. All cell culture plastics were from Sarstedt, unless otherwise specified. Plates were not coated prior to seeding cells. The absence of mycoplasma contamination was routinely confirmed by PCR using 5′-GGCGAATGGGTGAGTAACACG-3′ and 5′-CGGATAACGCTTGCGACTATG-3′ primers detecting 16S ribosomal RNA of mycoplasma in the media after 2–3 days of cell exposure.

### DNA constructs, cloning and mutagenesis

HiBiT-GPR183 plasmid DNA was generated with Gibson cloning using a codon-optimized GPR183 from GPR183-Tango plasmid DNA (#66342 Addgene, deposited by Bryan Roth) as an insert and HiBiT-FZD_6_ with a 5-HT_3A_ signal peptide plasmid DNA as a backbone [44]. In the newly generated construct, the GPR183 insert sequence replaced FZD_6_ insert sequence. In this construct, the N-terminally cloned HiBiT tag (GTGAGCGGCTGGCGGCTGTTCAAGAAGATTAGC) is followed by a GS linker (GGATCC, BamHI site). HiBiT-GPR183-Nluc construct was generated using Gibson cloning, inserting Nluc from Nluc-FZD_6_ [45] onto the C-terminus of HiBiT-GPR183, without a linker. HiBiT-GPR183-mNG-Nluc plasmid DNA constructs were generated using Gibson cloning, inserting mNeonGreen (mNG) from mNeonGreen-APEX plasmid DNA (#202591 Addgene, deposited by Reuben Harris) at three distinct positions into the GPR183 sequence (following P231, T233 or K235) in HiBiT-GPR183-Nluc. HA-GPR183-HiBiT was generated using HA-FZD_5_-HiBiT plasmid DNA and GPR183-Tango plasmid DNA. HA-GPR183-(T233)-mNG2_(11)_-HiBiT plasmid DNA was generated using the HA-GPR183-HiBiT and mNeonGreen2_(1-10)_/mNeonGreen2_(11)_ plasmid DNA (#82611 Addgene, deposited by Bo Huang). HiBiT-GPR183-Halo-Nluc plasmid DNA construct was generated using Gibson cloning, inserting Halo from FZD_5_-Halo-Nluc sensor ([46]) in HiBiT-GPR183-Nluc. Plasmid DNA constructs encoding different receptor mutants were generated with a GeneArt Site Directed mutagenesis kit (Thermo Fisher Scientific). rGFP-CAAX plasmid DNA, Rap1Gap-*R*luc2 plasmid DNA [47] and β-arrestin2-*R*luc2 in the pcDNA3.1(+) backbone were synthesized by GenScript. β-arrestin2-mVenus, HA-FZD_5_-HiBiT and LgBiT-CAAX [48] were kind gifts from Lukas Grätz (University of Bonn) and Gunnar Schulte (Karolinska Institutet). Plasmid DNA encoding an alpha subunit of Gi_1_ was from cDNA.org. Salmon sperm DNA (ssDNA) was from Thermo Fisher Scientific. The constructs were validated by Sanger sequencing (Eurofins GATC).

### Ligands

7α,25-dihydroxycholesterol (GPR183 agonist) was from Sigma (#SML0541). NIBR189 (GPR183 inverse agonist) was from Tocris (#5203). The compounds were dissolved in DMSO at 5 mM or 10 mM concentrations, aliquoted and stored at -20°C. Each aliquot was used a maximum of three times. For experiments, 7α,25-dihydroxycholesterol was used dissolved in 0.1% or 1% final DMSO concentration (please see the figure legends).

Compounds selected from the VS and the subsequent similarity search screen were purchased from Mcule (www.mcule.com), dissolved in DMSO at 1-10 mM concentrations, and stored at -20°C. Each aliquot was used a maximum of five times. The list of compounds can be found in the **Figure S1** and **Figure S2.**

### Gi protein activation with enhanced bystander BRET

HEK293A cells were transiently transfected in suspension using polyethylenimine (PEI, Polysciences). To measure Gi protein activation at the cell membrane interface, a total of ca. 4 × 10^5^ cells were transfected in 1 ml with 200 ng of HiBiT-GPR183 plasmid DNA constructs, 300 ng of rGFP-CAAX plasmid DNA, 100 ng of an alpha subunit of Gi_1_ plasmid DNA, 40 ng of Rap1Gap-*R*luc2 plasmid DNA and 360 ng of ssDNA. Next, transfected cells (4 × 10^4^ cells in 100 μl) were seeded onto white 96-well cell culture plates. Twenty-four hours later, the cells were washed once with 200 µl of HBSS (HyClone). Next, 70-90 μl of HBSS (depending on the assay setup) was added to the wells, and subsequently, 10 μl of coelenterazine 400a (2.5 μM final concentration, Biosynth) was added. The plate was incubated for 10 min, which was followed by the addition in the ligand-stimulation setups (keeping the final total solution volume at 100 μl in each well). Next, *R*luc2 emission (donor, 360-440 nm, 100 ms integration time) and rGFP emission (acceptor, 505-575 nm, 100 ms integration time) were measured at 37°C (three measurements for baseline, 10-25 minutes with the agonist and/or compounds – see the figure legends for detailed description for respective experimental paradigms). The ebBRET ratios were defined as acceptor emission/donor emission. The measurements were performed using a Tecan Spark microplate reader.

### β-arrestin2 recruitment with a direct BRET and ebBRET

HEK293A cells were transiently transfected in suspension using PEI. To measure β-arrestin2 recruitment in a direct BRET paradigm, a total of ca. 4 × 10^5^ cells were transfected in 1 ml with 50 ng of HiBiT-GPR183-Nluc plasmid DNA, 250 ng of β-arrestin2-mVenus plasmid DNA, 700 ng of ssDNA. To measure β-arrestin2 recruitment to the cell membrane-expressed GPR183 in an ebBRET paradigm, a total of ca. 4 × 10^5^ cells were transfected in 1 ml with 200 ng of HiBiT-GPR183 plasmid DNA, 50 ng of β-arrestin2-*R*luc2 plasmid DNA, 300 ng of rGFP-CAAX plasmid DNA and 450 ng of ssDNA. Next, for both experimental paradigms, transfected cells (4 × 10^4^ cells in 100 μl) were seeded onto white 96-well cell culture plates. Twenty-four hours later, the cells were washed once with 200 µl of HBSS (HyClone). Next, 70 μl of HBSS were added to the wells, and subsequently, 10 μl of furimazine (1:1000 final concentration, Promega) were added. The plate was incubated for 10 min, which was followed by the addition in the ligand-stimulation setups (keeping the final total solution volume at 100 μl in each well). Next, for the direct BRET setup, Nluc emission (donor, 460-500 nm, 50 ms integration time) and Venus emission (acceptor, 520-560 nm, 50 ms integration time) or, for the ebBRET setup, *R*luc2 emission (donor, 360-440 nm, 100 ms integration time) and rGFP emission (acceptor, 505-575 nm, 100 ms integration time), were measured at 37°C (three measurements for baseline, 10 min with the inverse agonists and then 15 minutes with the agonist). The BRET ratios were defined as acceptor emission/donor emission. The measurements were performed using a Tecan Spark microplate reader.

### GPR183 conformational sensors experiments

HEK293A cells were transiently transfected in suspension using PEI. To measure ligand-induced conformational change in GPR183 a total of ca. 4 × 10^5^ cells were transfected in 1 ml with 50 ng of HiBiT-GPR183-mNG-Nluc plasmid DNA constructs and 950 ng of ssDNA or 10 ng of HiBiT-GPR183-Halo-Nluc plasmid DNA and 990 ng of ssDNA. Next, transfected cells (4 × 10^4^ cells in 100 μl) were seeded onto white or black 96-well cell culture plates. Twenty-four to forty-eight hours later, the cells were washed once with 200 µl of HBSS (HyClone). Next, 70-80 μl of HBSS was added to the wells, and subsequently, 10 μl of 1:1000 final Nano-Glo substrate/furimazine (Promega) was added for the mNG-Nluc sensors or 50 μl of medium was removed and replaced with 50 μl of 1:2000 final Halo618 substrate (Promega) in the Halo-Nluc setup; the Halo-Nluc plate was washed the following day once with 200 µl of HBSS (HyClone), and subsequently, 10 μl of 1:1000 final Nano-Glo substrate/furimazine (Promega) was added. The plate was incubated for 10 min, which was followed by the ligand(s) addition (keeping the final total solution volume at 100 μl in each well). Next, for the mNG-Nluc sensors, Nluc emission (donor, 460-500 nm, 50 ms integration time) and mNG emission (acceptor, 505-545 nm, 50 ms integration time), and for the Halo-Nluc sensor, Nluc emission (donor, 445-470 nm, 50 ms integration time) and Halo618 emission (acceptor, 595-650 nm, 50 ms integration time) were measured at 37°C (three measurements for baseline, 10-25 minutes with the agonist and/or compounds – see the figure legends for detailed description for respective experimental paradigms). The BRET ratios were defined as acceptor emission/donor emission. The Z-factor was calculated as in [49] using 3 μM of 7**α**,25-dihydroxycholesterol-treated cells as positive control and 0.1% DMSO-treated cells as negative control. The measurements were performed using a Tecan Spark microplate reader.

### Generation of HEK293A cells stably-overexpressing HiBiT-GPR183-(T233)-mNG-Nluc

HEK293A cells stably overexpressing HiBiT-GPR183-(T233)-mNG-Nluc were generated following transfection of ca. 4 × 10^5^ cells with 1000 ng of HiBiT-GPR183-(T233)-mNG-Nluc plasmid DNA construct. The cells were seeded onto a 6-well plate (8 x 10^5^ cells) and cultured in DMEM (HyClone) supplemented with 10% fetal bovine serum. About 24 hours after the transfection, the cells were passaged at 1:10 and 24 hours later the cell medium was supplemented with 1000 µg/ml G418 (Thermo Fisher Scientific). The medium was replaced every two to three days to select the cells transfected with the plasmid. The cells were maintained in the presence of the antibiotic for a period of 31 days until a stable culture was established. The stability of protein expression was verified by measuring Nluc luminescence. The stable cell lines were maintained in complete DMEM medium in the presence of 250 µg/ml G418. The cell line was used only in the experiments to determine the Z-factor.

### Bystander BRET to measure the cell surface expression of C-terminally HiBiT-tagged receptors

HEK293A cells at a density of 4 x 10^5^ cells/ml were transfected in suspension using PEI with 500 ng of the indicated receptor plasmid DNA and 500 ng of LgBiT-CAAX plasmid DNA. The cells in a volume of 100 µl were seeded onto a white 96-well plate with a flat bottom. Twenty-four hours later, the cells were washed once with 200 µl of HBSS (HyClone). Next, 90 μl of HBSS were added to the wells, and subsequently, 10 μl of furimazine (1:1000 final concentration, Promega) were added. The plate was incubated for 10 minutes and subsequently Nluc/NanoBiT luminescence (460-500 nm, 50 ms integration time) was measured using a Tecan Spark microplate reader.

### In vitro cell migration assay

*In vitro* cell migration was assessed using 96-well HTS transwell plates with an 8 µm pore size (Corning). Briefly, peripheral blood mononuclear cells (PBMC) from healthy donors (n=6) (500,000 cells/well) were stimulated with 10 µM 7a,25-dihydroxycholesterol in serum-free RPMI. After 4 hours incubation at 37°C with 5% CO_2_, cells that had migrated to the lower chamber were harvested. They were first incubated with a viability dye for 10 mins at 4°C and then stained with fluorescently labelled antibodies, as shown in supplementary **Table S1**, for 20 mins at 4°C. RANTES (CCL5; 50 ng/ml) and unstimulated cells (US) were used as a positive and negative controls, respectively. To inhibit GPR183-mediated migration, PBMC were pre-incubated with 10 µM of compound **78** or NIBR189 at 37°C for 30 mins. The cells were then transferred to the transwell plate and processed as mentioned above. Flow cytometry analysis was performed using FlowJo version 10.10.

### Statistical analysis

All data presented in the main figures come from at least three individual experiments (biological replicates), with each individual experiment typically performed at least in duplicates (technical replicates) for each condition, unless otherwise specified in a figure legend. One biological replicate refers to wells containing cells seeded from the same individual cell culture flasks and measured on the same day. Different biological replicates were transfected using separate transfection mixtures and measured on different days. Technical replicates are defined as individual wells with cells from the same biological replicate. Data presented in the Supporting Information, which indeed play a supporting role, may come from fewer than three biological experiments, and furthermore can be presented in the form of a representative example, therefore without a statistical analysis. Samples were not randomized or blinded during the experiments. Statistical and graphical analyses were performed using Graph Pad Prism software. Two datasets were analysed for statistical differences with paired *t*-test or Mann-Whitney test. Three or more datasets were analysed by one-way ANOVA with multiple comparison Dunnett’s post-hoc analysis or Kruskal-Wallis with multiple comparison Dunn’s post-hoc analysis. Significance levels are given and displayed in the figures as: *P < 0.05; **P < 0.01; ***P < 0.001; ****P < 0.0001. Differences between datasets which did not reach statistical significance are left unmarked or marked with “ns”. Data points throughout the manuscript are indicated as the mean ± standard error of the mean (s.e.m.) unless otherwise stated.

## RESULTS

*AI-enforced VS.* To discover novel GPR183 binders, the PyRMD2Dock protocol was employed as depicted in **Figure 1**. It combines the Ligand-Based VS (LBVS) tool PyRMD [50] with one of the most widely used docking software, AutoDock-GPU (AD4-GPU) [34]. By implementing PyRMD2Dock, it is possible to rapidly screen large chemical databases and identify those with the highest predicted binding affinity to a target protein. The first step of this workflow was to randomly select 1M compounds from the screened database (in our case we selected a set of ∼10M ligands from a subset of lead-like purchasable ZINC20 database [51]), and subject them to docking calculations employing the solved Cryo-EM structure of GPR183 in complex with its endogenous ligand 7a,25-dihydroxycholesterol (PDB 7TUZ [2]). For all the compounds, the predicted lowest binding free energy (Δ*G*_AD4_) was extracted, and the results were analysed based on their Δ*G*_AD4_ value distribution. In this study, compounds were categorized as “active” or “inactive” based on their docking scores (ΔG_AD4_). To establish these classes, we defined five distinct “activity” thresholds (≤ -9.5, -10.0, -10.5, -10.75, and -11.0 kcal/mol) and five “inactivity” thresholds (≥ - 4.0, -4.5, -5.0, -5.5, and -6.0 kcal/mol). This created 25 unique datasets by combining each active threshold with each inactive threshold. The performance of our classification model is highly dependent on the ε cutoff values, which define the maximum allowed projection distance for active and inactive compounds within the training’s linear subspace. To optimize these parameters, we performed a comprehensive benchmarking analysis. For each of the 25 datasets, we employed a repeated stratified κ-fold cross-validation (5 folds, 3 repetitions) and systematically tested ε cutoff values from 0.01 to 0.99 (in steps of 0.10) for both classes. The optimal model was identified by the best trade-off between the True Positive Rate (TPR) and False Positive Rate (FPR). This was achieved using the dataset defined by an active threshold of ≤ -10.0 kcal/mol and an inactive threshold of ≥ -6.0 kcal/mol. The best-performing model, which yielded a TPR/FPR trade-off of 0.69, utilized an ε cutoff value of 0.99 for the active set and 0.80 for the inactive set. The remaining ∼9M compounds from the selected chemical databases were screened using the PyRMD screening module trained with the docking results, employing default settings and the ε cutoff values reported above. PyRMD returned a list of compounds classified as possible GPR183 binders with an associated confidence score (RMD score) of its predictions. Subsequently, the best 50 000 GPR183 binders predicted by PyRMD (i.e., the ones with the highest RMD score) were then docked into the GPR183 receptor, and to further refine the selection, we subsequently applied a pose filtering scheme aimed at selecting the ligands that were predicted to directly contact Y116^3.37^, Y260^6.51^, R87^2.60^, and Y112^3.33^ (numbers in the superscript indicate residue positions according to the Ballesteros-Weinstein numbering scheme [2, 52]). These selection criteria, and a subsequent visual inspection, led to the selection of 70 GPR183 candidate binders

**Figure 1.**
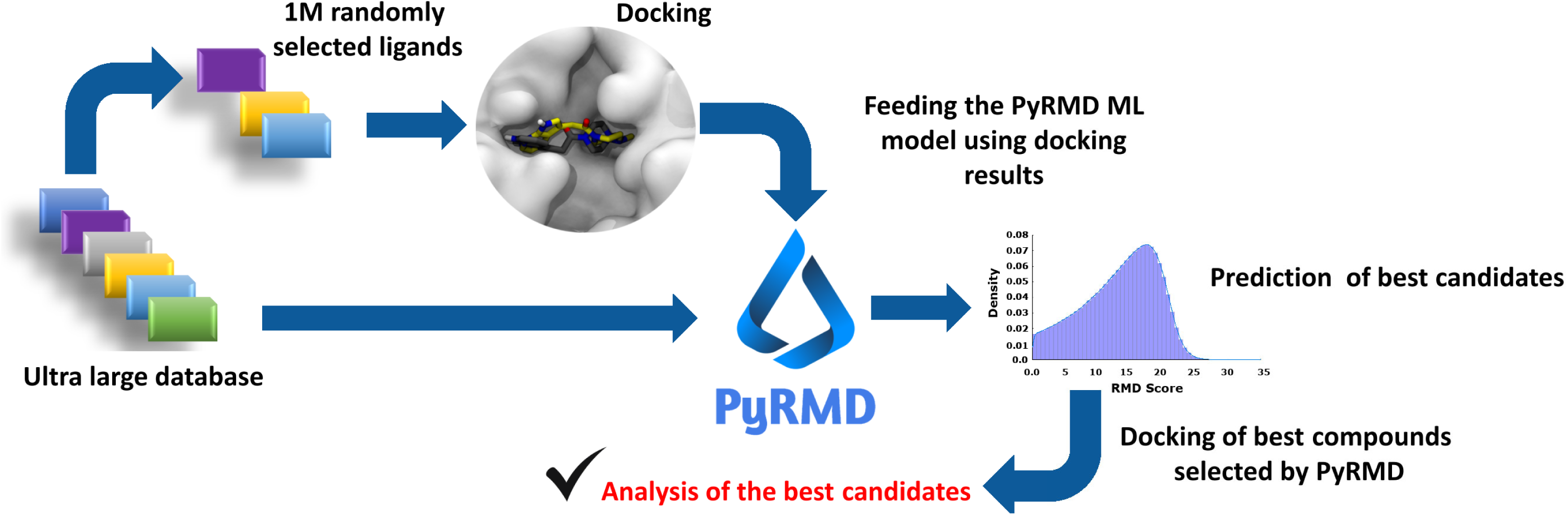
The PyRMD2Dock workflow.

### GPR183-mediated Gi activation – compounds from the AI-enforced VS

Based on the VS results, 70 compounds were purchased and analysed in ebBRET-based assays to measure their effects on the activation of Gi mediated by the constitutive activity of overexpressed GPR183, at the cell membrane. In the first set of experiments, we have evaluated all 70 compounds at a fixed concentration of 10 μM – the data can be found in **Figure 2a**. Using the cut-off defined by the statistically significant difference (one-way ANOVA test with Dunnett’s post-hoc analysis, P<0.05) in the ebBRET between the vehicle (DMSO)-treated cells and compound-treated cells, we selected two compounds that significantly reduced the constitutive activity: **3** (4-{2-[4-(2-fluorophenyl)-4-hydroxy-octahydro-1H-isoindol-2-yl]-2-oxoethyl}-1,2-dihydrophthalazin-1-one) and **43** (2-{[1-(naphthalene-2-carbonyl)piperidin-4-yl]methyl}-6-(1H-pyrazol-1-yl)-2,3-dihydropyridazin-3-one). Had a less stringent selection cut-off been defined using the LSD-Fisher post-hoc test, an additional 12 compounds would have met the criteria for significant reduction of the ebBRET signal (compounds: 4, 15, 20, 21, 35, 36, 37, 41, 42, 45, 51, 53). This cohort represents 20% of the total compounds evaluated in these assays, which is in line with the success rate expected in structure-based VS campaigns. The representative kinetic plots for this group can be found in **Figure S3**.

**Figure 2.**
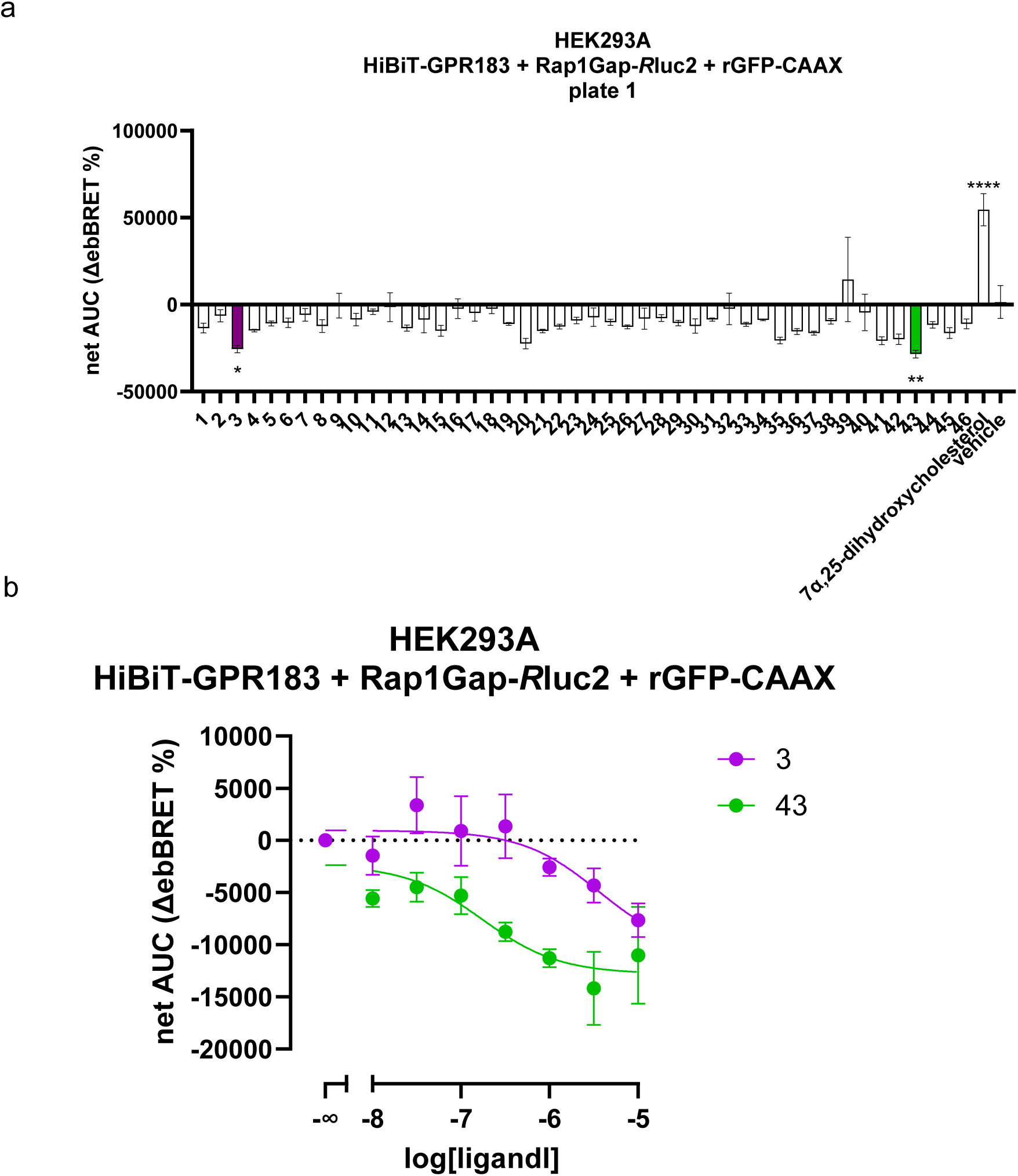
The compounds inhibit the constitutive activity of GPR183. **a)** The results from the 10 μM compounds screen using the ebBRET-based Gi activation assay with an overexpressed HiBiT-GPR183. The compounds were added to the cells and the ebBRET measured over 15 minutes. The data are presented as net AUC of ΔebBRET % over the three baseline reads measured prior to the compound addition; vehicle (0.1% DMSO) was not subtracted. The data are shown as mean ± s.e.m. of three independent experiments. The data were analysed for differences with ANOVA with Dunnett’s post-hoc analysis; *P < 0.05; **P < 0.01; ****P < 0.0001. The selected compounds, **3** and **43**, are shown as purple and green bars, respectively. **b)** The concentration response curve of compounds **3** and **43** in the same assay. The compounds were added, and the ebBRET signal was measured over 15 minutes. The data are presented as net AUC of ΔebBRET % over the three baseline reads measured prior to the compound addition; vehicle (0.1% DMSO) was subtracted. The data are shown as mean ± s.e.m. of three independent experiments.

Compounds **3** and **43** were then studied in the next step **(Figure 2b)**, where the aforementioned ebBRET-based assay was reiterated by applying a range of different concentrations of the two molecules, spanning from 10 nM to 10 μM. In these assays, compound **3** inhibited the GPR183-mediated Gi activation with p*IC*_50_ = 5.44 (95% CI: not converged – 6.44), and compound **43** inhibited the GPR183-mediated Gi activation with a higher potency and p*IC*_50_ = 6.76 (95% CI: 5.93 – 7.86). To this end, these data supported the selection of compound **43** for further investigations.

### Hit expansion and GPR183-mediated Gi activation and β-arrestin2 recruitment

To further explore the chemical space around this promising hit, a similarity search was performed to identify derivatives of compound **43**. This involved selecting compounds featuring structural similarity with the hit (Tanimoto index > 0.70). Through this process, we identified 34 potential derivatives (**Table 1)** that were purchased and assessed in the same ebBRET-based Gi activation experimental setup at a fixed concentration of 10 μM – the data can be found in **Figure 3a** and **Table 1**.

**Figure 3.**
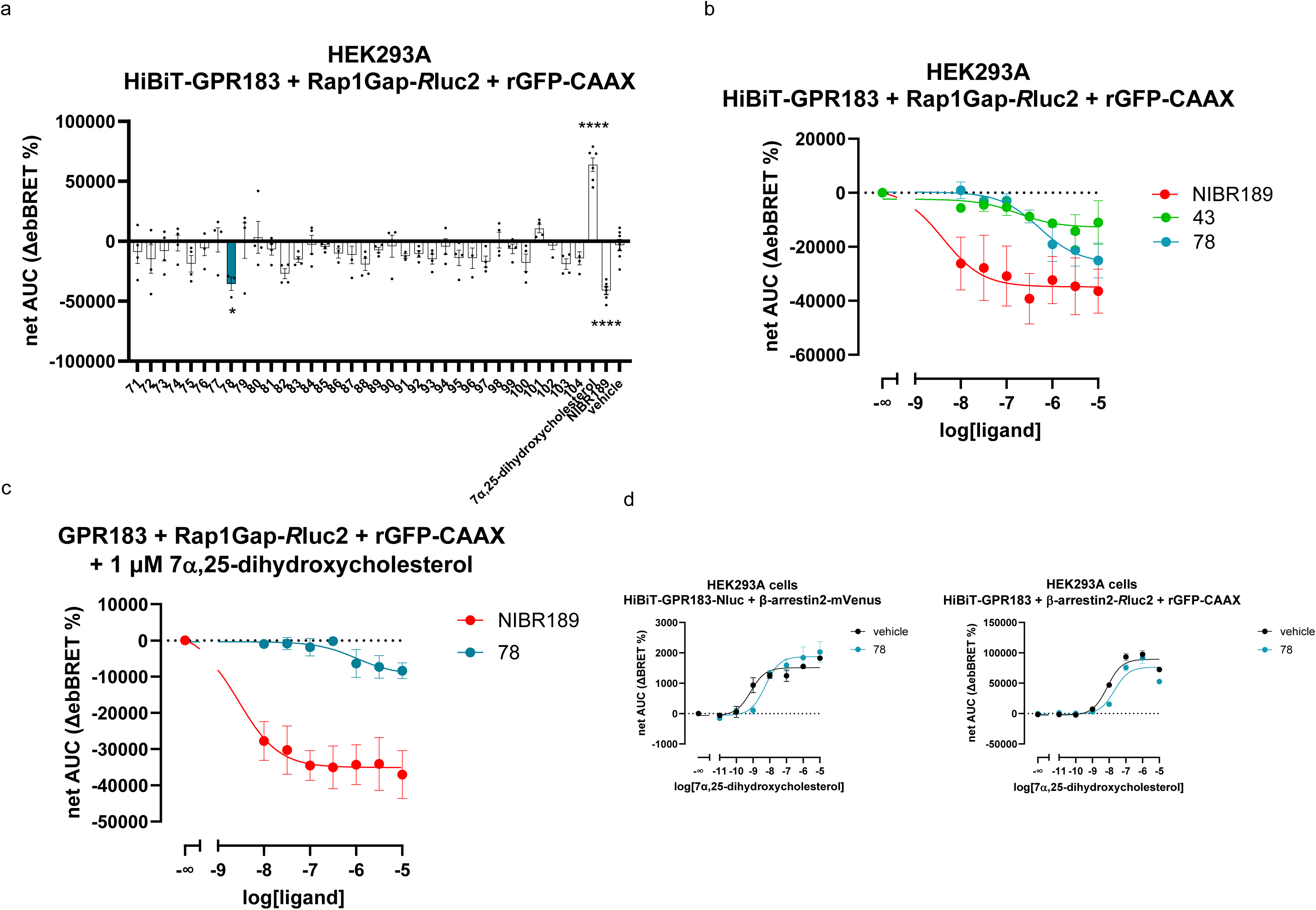
Analogues of compound 43 inhibit constitutive and agonist-stimulated GPR183 activity. **a)** The results from the screen with the 34 analogues of **43.** The compounds were used at 10 μM and their activity to modulate HiBiT-GPR183 activation assesses using the ebBRET-based Gi activation assay. The compounds were added to the cells and the ebBRET measured over 15 minutes. The data are presented as net AUC of ΔebBRET % over the three baseline reads measured prior to the compound addition; vehicle (0.1% DMSO) was not subtracted. The data are shown as mean ± s.e.m. of three-four independent experiments. The data were analysed for differences with ANOVA with Dunnett’s post-hoc analysis; *P < 0.05; **P < 0.01. The selected compound **78** is shown in dark green bars. **b)** The concentration response curve of compounds **43**, **78** and NIBR189 in the same assay. The compounds were added, and the ebBRET signal was measured over 15 minutes. The data are presented as net AUC of ΔebBRET % over the three baseline reads measured prior to the compound addition; vehicle (0.1% DMSO) was subtracted. The data are shown as mean ± s.e.m. of three independent experiments. The same data for the **43** are also shown in the Figure 2b. **c)** The concentration response curve of compounds **78** and NIBR189 following cells incubation with 1 μM 7α,25-dihydroxycholesterol for 5 minutes. The data are presented as net AUC of ΔebBRET % over the three baseline reads measured prior to the compounds’ addition; vehicle (0.1% DMSO) was subtracted. The data are shown as mean ± s.e.m. of three-four independent experiments. **d)** Agonist-stimulated HiBiT-GPR183-Nluc- or HiBiT-GPR183-mediated recruitment of β-arrestin2-mVenus (**left**) and β-arrestin2-*R*luc2 (**right**). The concentration response curve of 7α,25-dihydroxycholesterol in the presence or absence (vehicle, 1% DMSO) of 10 μM of **78**. **78** or vehicle were added, and the BRET/ebBRET signal was measured over 10 minutes, followed by the addition of different agonist concentrations. The plate was measured for another 15 minutes. The data are presented as net AUC of ΔBRET %/ΔebBRET over the first 10-minute-long read measured prior to the agonist addition; vehicle (1% DMSO) was subtracted. The data are shown as mean ± s.e.m. of three independent experiments. The difference in agonist potency was assessed by comparing log*EC*_50_ values for the two curves with the extra sum-of-squares F test; P = 0.0048.

**Table 1.**
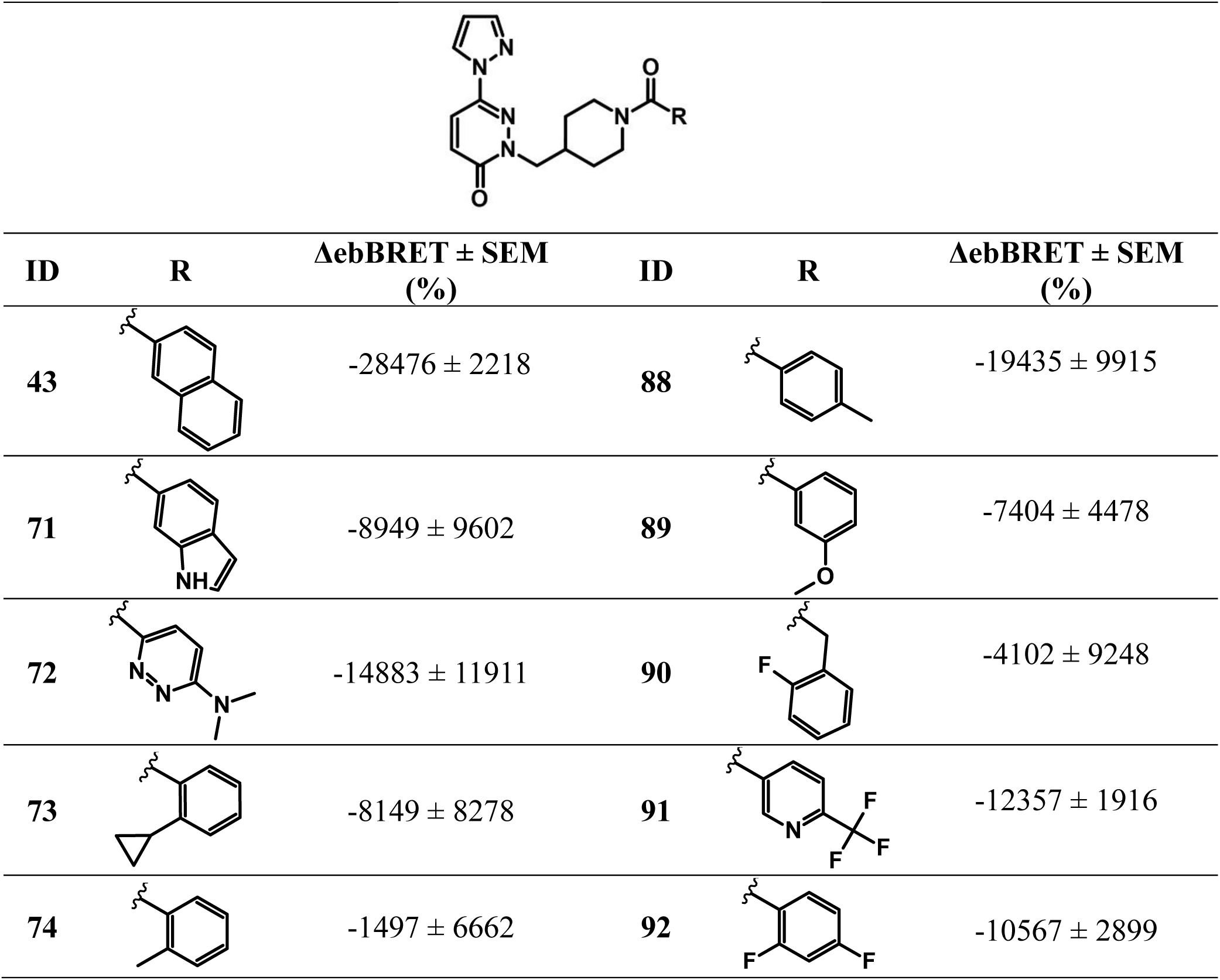
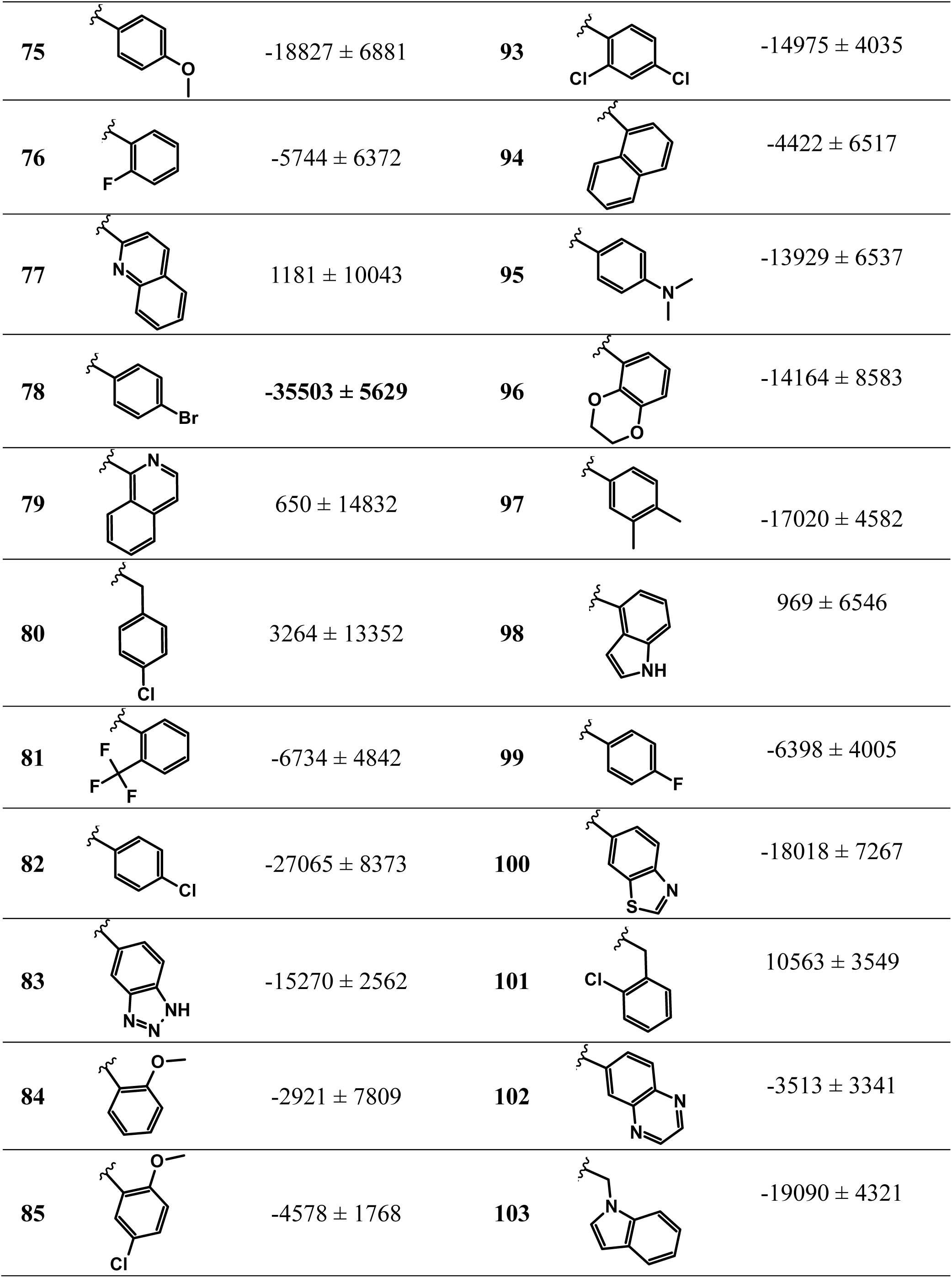
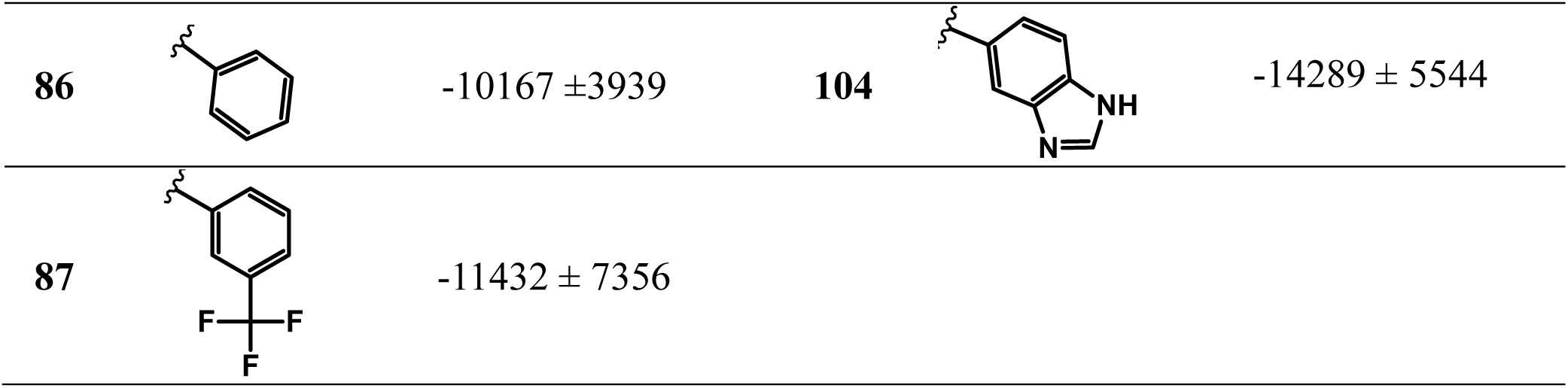
Structures of the 34 analogues of 43, along with their experimental ΔebBRET (%).

|  |  |  |  |  |  |
| --- | --- | --- | --- | --- | --- |
| ID | R | $\Delta\text{ebBRET} \pm \text{SEM}$<br>(%) | ID | R | $\Delta\text{ebBRET} \pm \text{SEM}$<br>(%) |
| 43 | | $-28476 \pm 2218$ | 88 | | $-19435 \pm 9915$ |
| 71 | | $-8949 \pm 9602$ | 89 | | $-7404 \pm 4478$ |
| 72 | | $-14883 \pm 11911$ | 90 | | $-4102 \pm 9248$ |
| 73 | | $-8149 \pm 8278$ | 91 | | $-12357 \pm 1916$ |
| 74 | | $-1497 \pm 6662$ | 92 | | $-10567 \pm 2899$ |
| 75 | | $-18827 \pm 6881$ | 93 | | $-14975 \pm 4035$ |
| 76 | | $-5744 \pm 6372$ | 94 | | $-4422 \pm 6517$ |
| 77 | | $1181 \pm 10043$ | 95 | | $-13929 \pm 6537$ |
| 78 | | $-35503 \pm 5629$ | 96 | | $-14164 \pm 8583$ |
| 79 | | $650 \pm 14832$ | 97 | | $-17020 \pm 4582$ |
| 80 | | $3264 \pm 13352$ | 98 | | $969 \pm 6546$ |
| 81 | | $-6734 \pm 4842$ | 99 | | $-6398 \pm 4005$ |
| 82 | | $-27065 \pm 8373$ | 100 | | $-18018 \pm 7267$ |
| 83 | | $-15270 \pm 2562$ | 101 | | $10563 \pm 3549$ |
| 84 | | $-2921 \pm 7809$ | 102 | | $-3513 \pm 3341$ |
| 85 | | $-4578 \pm 1768$ | 103 | | $-19090 \pm 4321$ |
| 86 | | $-10167 \pm 3939$ | 104 | | $-14289 \pm 5544$ |
| 87 | | $-11432 \pm 7356$ | | | |

Again, using the cut-off defined by the statistically significant difference (one-way ANOVA test with Dunnett’s post-hoc analysis, P<0.05) in the ebBRET between the vehicle (DMSO)-treated cells and compound-treated cells, we have selected compound **78** (2-{[1-(4-bromobenzoyl)piperidin-4-yl]methyl}5.-6-(1H-pyrazol-1-yl)-2,3-dihydropyridazin-3-one). If a less stringent LSD-Fisher post-hoc test had been used to define a selection cut-off, one more compound would have significantly reduced the ebBRET signal (compound 82), equalling 5.9 % of the total number of compounds tested in these assays.

In the second step, we proceeded with compound **78** and performed the aforementioned ebBRET-based assay applying a range of different concentrations of the molecule, spanning from 10 nM to 10 μM. In these assays, compound **78** inhibited the GPR183-mediated Gi activation with p*IC*_50_ = 6.29 (95% CI: 5.69 – 6.86). The potency was no different in comparison with the parent compound **43** (extra sum-of-squares F test P=0.43), but the efficacy (ebBRET_min_) was higher for compound **78** (extra sum-of-squares F test P=0.01) (**Figure 3b**). NIBR189 was used as a positive control in the assay. We confirmed a high efficacy and potency of NIBR189 as an inverse agonist (p*IC*_50_=8.39 (95% CI: 7.90 – 9-66)).

Next, we also used the **78** in a competition assay where the compound was incubated together with the agonist 7α,25-dihydroxycholesterol (1.0 μM). Following a set of 15-minute-long kinetic experiments, we established that the **78** inhibited the agonist-induced GPR183-mediated activation of Gi with the p*IC*_50_=5.99 (95% CI: 5.16 – 6.70). Again, NIBR189, (p*IC*_50_=8.54 (95% CI: 8.30 – 8.91)), was used as a positive control in the “competition-mode” assay (**Figure 3c**).

Finally, we assessed the ability of **78** to inhibit recruitment of β-arrestin2 to the agonist-stimulated receptor in two experimental setups, assessing 1) direct BRET between the receptor and β-arrestin2 (Nluc-Venus) as well as using 2) an indirect, ebBRET setup that measures signal between the β-arrestin2 and membrane-anchored CAAX (*R*luc2-rGFP), which indicates the recruitment of this scaffold protein to agonist-activated cell membrane-present GPR183. Here, we found that **78** significantly reduced the potency of the agonist to promote β-arrestin2 recruitment in two independent experimental paradigms (extra sum-of-squares F test, P = 0.0048 and P = 0.038, respectively; **Figure 3d**). The consistency of this effect across distinct assay formats and BRET pair combinations indicates that the observed inhibition is robust and unlikely to be an assay-specific artifact. Taken together with previous data demonstrating Gi protein activation, these results support the conclusion that **78** acts as a balanced ligand at GPR183.

### GPR183 conformational sensors

To analyse the binding of the **78** to GPR183 in greater detail, we have generated a novel intramolecular conformational sensor based on BRET between mNG located in the ICL3 and Nluc fused to the C-terminus (**Figure 4a**). First, we screened three potential insertion sites for the mNG in the ICL3: P231, T233, and K235. While all three sensors responded to the agonist stimulation by a similar reduction in BRET (**Figure 4b**) and could be in principle used in the pharmacological validation of the binding compounds, the T233 sensor had the biggest apparent signal window and a relatively stable temporal resolution, thus, this HiBiT-GPR183-(T233)-mNG-Nluc construct was selected for further investigations. This sensor variant exhibited an acceptable Z-factor = 0.30 ± 0.13 (**Figure S4a**). Next, the introduction of the mNG tag into the ICL3 likely compromises the constitutive and ligand-stimulated ability of GPR183 to activate Gi as shown for an ebBRET Gi assay-compatible HA-GPR183-(T233)-mNG2_(11)_-HiBiT sensor in which the ICL3 was modified with a much smaller tag (**Figure S4b**). On the other hand, a corresponding to HiBiT-GPR183-(T233)-mNG-Nluc construct sensor based on Halo and Nluc did not respond to the agonist stimulation (**Figure S4c**). Next, we used the HiBiT-GPR183-(T233)-mNG-Nluc sensor to analyse the half-maximal response of the agonist 7α,25-dihydroxycholesterol (**Figure 4c**). Given that the conformational response of the receptor is the first direct and very proximal event that follows an agonist binding, we interpret that for this experimental setup the agonist *EC*_50_ ≈ *K*_d_. To this end, we measured the p*K*_d_ = 9.15 (95% CI = 8.75 – 9.56). The addition of different concentrations of NIBR189 leads to a concentration-dependent inhibition of the agonist-induced change in BRET (**Figure 4d**) with a p*K*_i_ = 9.83 (95% CI = 9.19 – 10.75).

**Figure 4.**
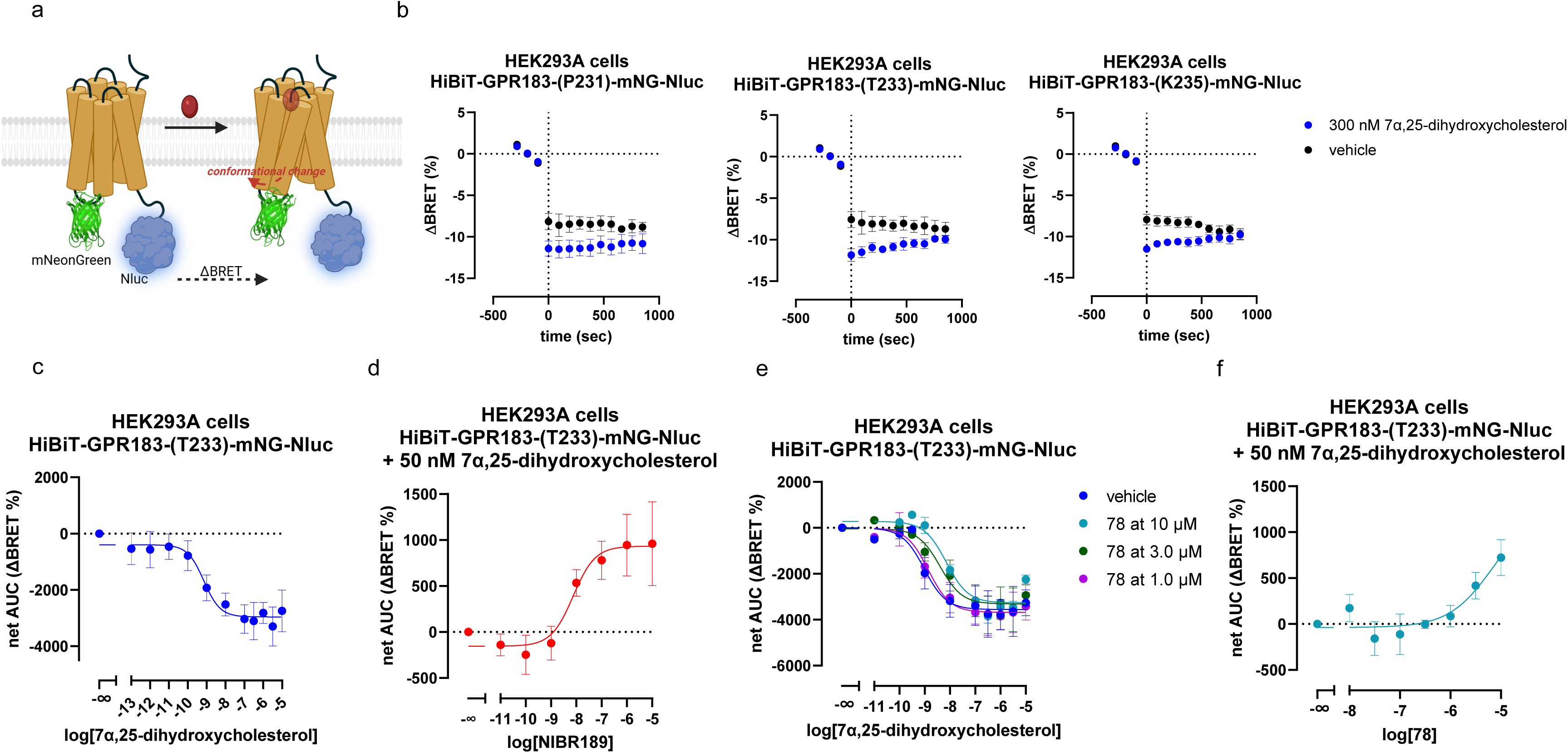
The novel GPR183 conformational sensor allows pharmacological profiling of receptor ligands. **a)** The schematic representation on the HiBiT-GPR183-mNG-Nluc conformational sensor. The mNeonGreen (mNG) is inserted into the intracellular loop 3 (ICL3) and the Nanoluciferase (Nluc) is fused directly to the C-terminus. **b)** Agonist stimulation of the three generated sensors, in which the mNG tag was inserted directly after P231, T233 or K235, leads to a reduction in BRET. The data are presented as 15-min kinetic plots with ΔBRET % (over the three baseline reads); vehicle (1% DMSO) was not subtracted. The data are shown as mean ± s.e.m. of three independent experiments**. c)** The concentration response curve of the agonist. The data are presented as net AUC of ΔBRET % over the three baselines reads measured prior to the agonist addition; vehicle (1.0% DMSO) was subtracted. The plate was measured for another 15 minutes. The data are shown as mean ± s.e.m. of five independent experiments. **d)** The concentration response curve of the inverse agonist NIBR189. Different concentrations of NIBR189 were added, and the BRET signal was measured over 10 minutes, followed by the addition of 50 nM 7α,25-dihydroxycholesterol. The plate was measured for another 15 minutes. The data are presented as net AUC of ΔBRET % (over the first 10-minute-long read); vehicle (1% DMSO) was subtracted. The data are shown as mean ± s.e.m. of three independent experiments. **e)** The concentration response curve of 7α,25-dihydroxycholesterol in the presence or absence (vehicle) of 10 μM, 3.0 μM or 1.0 μM of **78**. **78** or vehicle (0.1% DMSO) were added and the plate incubated for 10 minutes at 37°C. The three baseline measurements were read and they were followed by the addition of different agonist concentrations and the plate was measured for 15 minutes. The data are presented as net AUC of ΔBRET % (over the three baseline measurements); vehicle (1% DMSO) was subtracted. The data are shown as mean ± s.e.m. of four to five independent experiments. The data were analysed using the Gaddum/Schild EC50 shift function in GraphPad Prism. **f)** The concentration response curve of **78**. Different concentrations of **78** were added, and the BRET signal was measured over 10 minutes, followed by the addition of 50 nM 7α,25-dihydroxycholesterol. The plate was measured for another 15 minutes. The data are presented as net AUC of ΔBRET % (over the first 10-minute-long read); vehicle (1.0% DMSO) was subtracted. The data are shown as mean ± s.e.m. of three independent experiments.

Employing the HiBiT-GPR183-(T233)-mNG-Nluc sensor, we then analysed the binding of the **78** and reported that it right-shifted the agonist concentration curve with a Gaddum/Schild analysis, with the Schild slope =1, indicating the p*K***_b_** = 5.66 (95% CI: 4.91 – 6.26; **Figure 4e)**. In a setup, where a full concentration range of **78** was used in the presence of 50 nM 7α,25-dihydroxycholesterol, the p*K*_i_ = 7.02 (95% CI = 6.08 – 7.95), however, as expected, the curve did not reach saturation at the highest applied, 10 μM concentration of the compound **78**, and thus the parameter represents an apparent value calculated with a quite large confidence interval (**Figure 4f**).

### Molecular dynamics simulation and in vitro validation of the compound **78** binding site

To provide a picture from an atomistic point of view of the binding properties of **78** in complex with GPR183 protein, *in silico* studies were undertaken employing AD4-GPU docking software [34]. As expected, the *in silico* predicted binding mode of **78** closely resembles that of the experimentally determined 7α,25-dihydroxycholesterol (**Figure 5a**). From an overall perspective, **78** is located deep within the central cavity of the GPR183 receptor’s transmembrane domain, adopting an elongated conformation that allows it to establish contacts with TM3 through TM7 (**Figure 5b**). Specifically, the *p-*bromophenyl moiety forms extended π-π interactions with the nearby Y116^3.37^ (Ballesteros-Weinstein numbering scheme [52]) side chain located on TM3 (**Figure 5c**). This interaction is strengthened by the electron-withdrawing bromine substituent on the phenyl ring, which reduces the electron density of the π-cloud, thereby promoting a robust π-stacking interaction [53]. In the predicted binding mode of **78**, the pyridazinone core establishes proficient H-bond interactions with the Y260^6.51^ side chain while the pyrazole moiety H-bonds both Y260^6.51^ and Y112^3.33^ side chains (**Figure 5c**).

**Figure 5.**
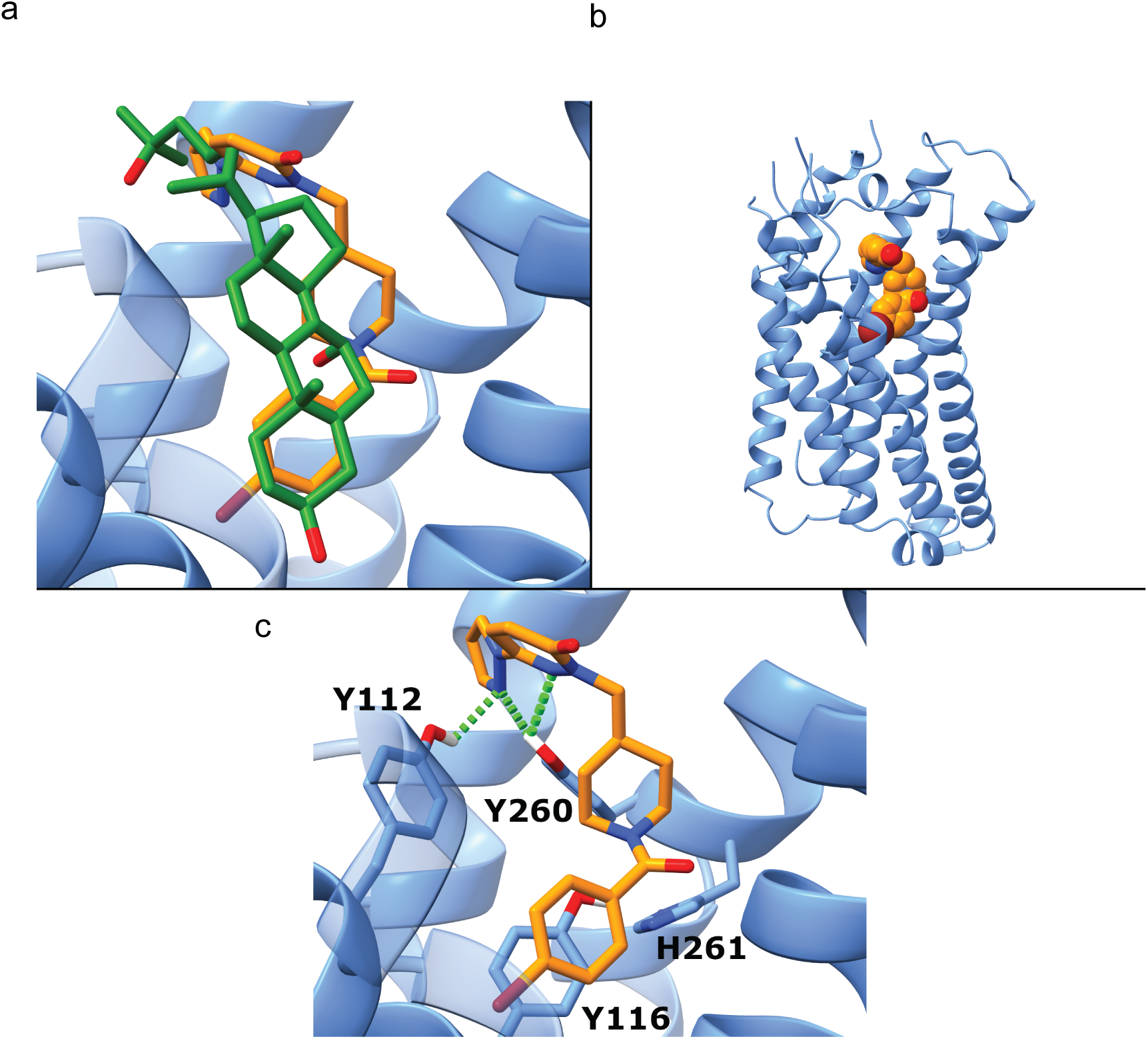
The binding pose of 78 in the transmembrane core of GPR183. **a)** Overlap between **78** and 7α,25-dihydroxycholesterol. Compound **78** is represented as orange sticks, and 7α,25-dihydroxycholesterol as green sticks, while the GPR183 protein is shown as blue ribbons and sticks. **b)** Overview of **78** binding mode. The ligand is depicted as orange spheres. **c)** Predicted binding mode of **78** in complex with GPR183 protein. H-bonds are represented as green dashes

To assess if the binding pose can be maintained, a 500 ns molecular dynamics (MD) simulation was performed on **78** in complex with GPR183 receptor using Amber22 software (Amber manual, Amber 2025, University of California, San Francisco). Analysis of the ligand root mean square deviation (RMSD) (**Figure 6a**) demonstrated that the pose of **78** was stable over the full 500 ns MD simulation time, exhibiting an average RMSD value of 0.51 Å. Subsequently, to further inspect the stability of the π-π interactions between **78** and the nearby Y116^3.37^ side chain, an analysis of the distances between the centres of mass of the *p*-bromophenyl moiety of **78** and the Y116^3.37^ side chain was performed, and the angles between the normal vectors of the two planes (delineated by the *p*-bromophenyl moiety of **78** and the Y116^3.37^ side chain, respectively) were also analysed (**Figure 6 b,c**). This comprehensive analysis further predicted the stability of the predicted π-π interaction, revealing an average distance between the planes of 3.77 Å and an average angle between their normal vectors of ±11.88°.

**Figure 6.**
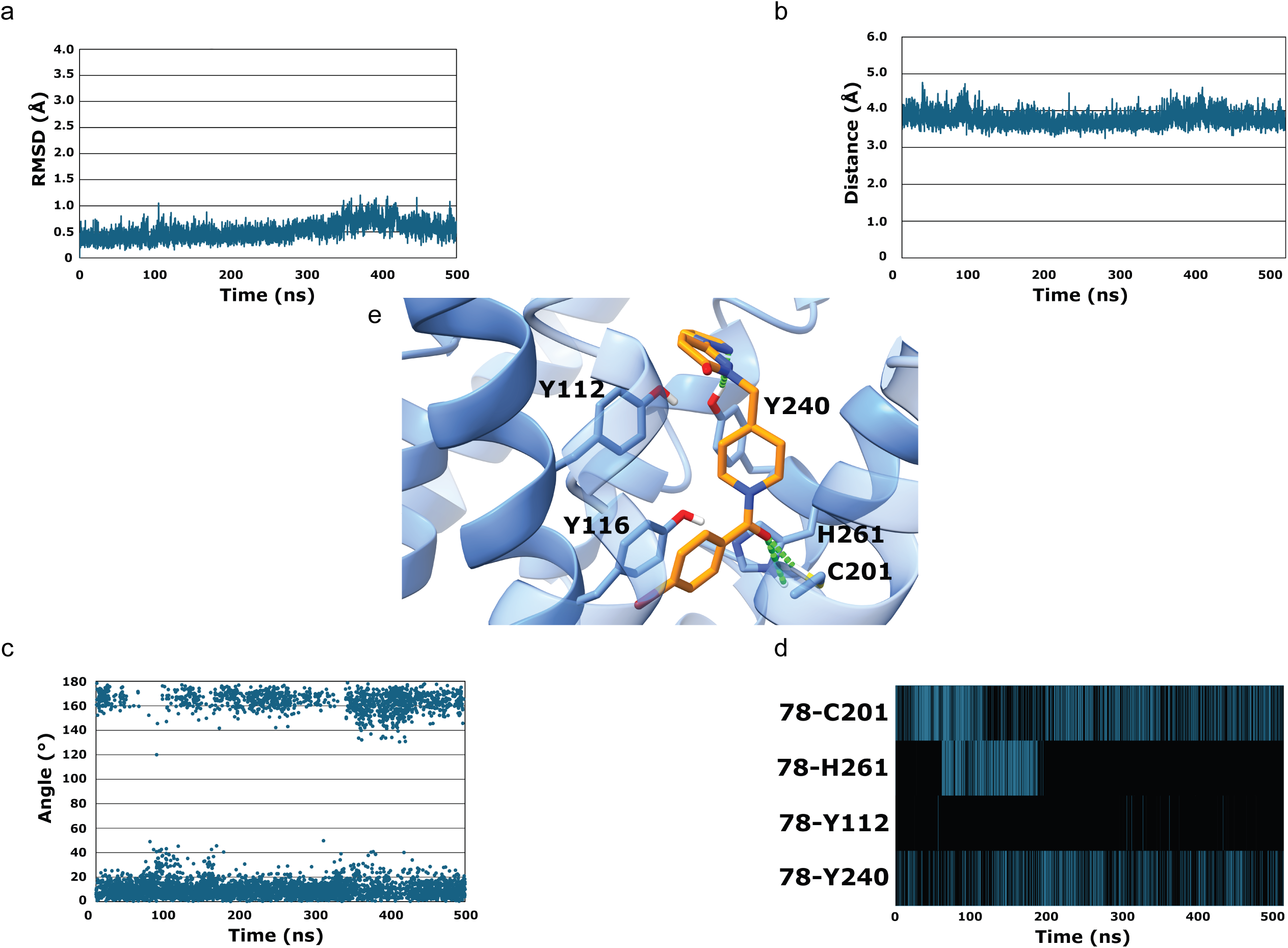
The key interactome of 78 in the GPR183: MD analysis. RMSD **(a)**, distance **(b)**, and angles **(c)** plots over time (in nanoseconds) of compound **78**. **d)** H-bond interactions vs trajectory time between **78** and C201, H261, Y112, and Y240. **e)** Representative structure of the single, well-defined cluster of **78** in complex with GPR183 protein. The ligand is depicted as orange sticks while the protein is shown as blue sticks and ribbons. H-bond are represented as green dashes.

Subsequently, analysis of the H-bonds between **78** and the GPR183 protein was conducted throughout the entire simulation. This analysis unveiled a strong and stable H-bond interaction between **78** and Y260^6.51^ (**Figure 6d**). In the same context, while an initial H-bond interaction between **78** and the Y112^3.33^ side chain was observed in the predicted binding mode, it proved to be unstable during the MD simulation. It may be tempting to postulate that this instability is ascribable to a minor rearrangement of the ligand molecule, evident in the change in geometry of its pyrazole ring (**Figure 6e**). As the pyrazole ring rotated, it lost the ability to interact with the Y112^3.33^ side chain. However, a detailed analysis of the interactions during the 500 ns of molecular dynamics allowed for the identification of two additional H-bond interactions between **78** and C201^5.43^ and H261^6.52^ side chains (**Figure 6e**).

To experimentally validate the binding mode predicted by our MD studies, we sought to confirm the importance of the key hydrogen bonding residues. We hypothesized that if Y260^6.51^ and H261^6.52^ are essential for binding of **78**, then mutating these residues should alter the compound’s ability to inactivate GPR183 and/or compete with 7α,25-dihydroxycholesterol. To test this, we engineered two receptor mutants, Y260F^6.51^ and H261A^6.52^, and subjected them to experiments assessing basal and agonist-induced receptor-mediated Gi activation as well as agonist-induced conformational change in the receptor.

In the case of the H261A^6.52^ mutant, 10 µM of compound **78** failed to alter the already very weak, in comparison with the GPR183 wild-type (**Figure 7a**), constitutive activity of GPR183 in the Gi activation assay (**Figure 7b**). Given the low basal activity of this mutant, we sought to confirm the apparent lack of compound **78** activity in this experimental paradigm by pre-applying the compound prior to the agonist stimulation. No significant difference was observed in agonist potency between vehicle and 10 µM **78**-treated conditions (p*EC*₅₀ = 7.04 (95% CI: 6.48–7.81) vs. p*EC*₅₀ = 7.33 (95% CI: 6.95–7.72); P = 0.43; **Figure 7b**). To further validate these findings, we evaluated whether 10 µM of **78** could suppress agonist-induced conformational changes in the GPR183 H261A^6.52^ mutant using the GPR183-(T233)-mNG-Nluc sensor. Consistent with the functional Gi-based assay, **78** did not significantly inhibit agonist activity/affinity in this context (p*K*_d_ = 6.90 (95% CI: 4.45–8.23) vs. p*K*_d_ = 7.42 (95% CI: 5.83–11.75) for vehicle and **78**-treated conditions, respectively; P = 0.55; **Figure 7c**).

**Figure 7.**
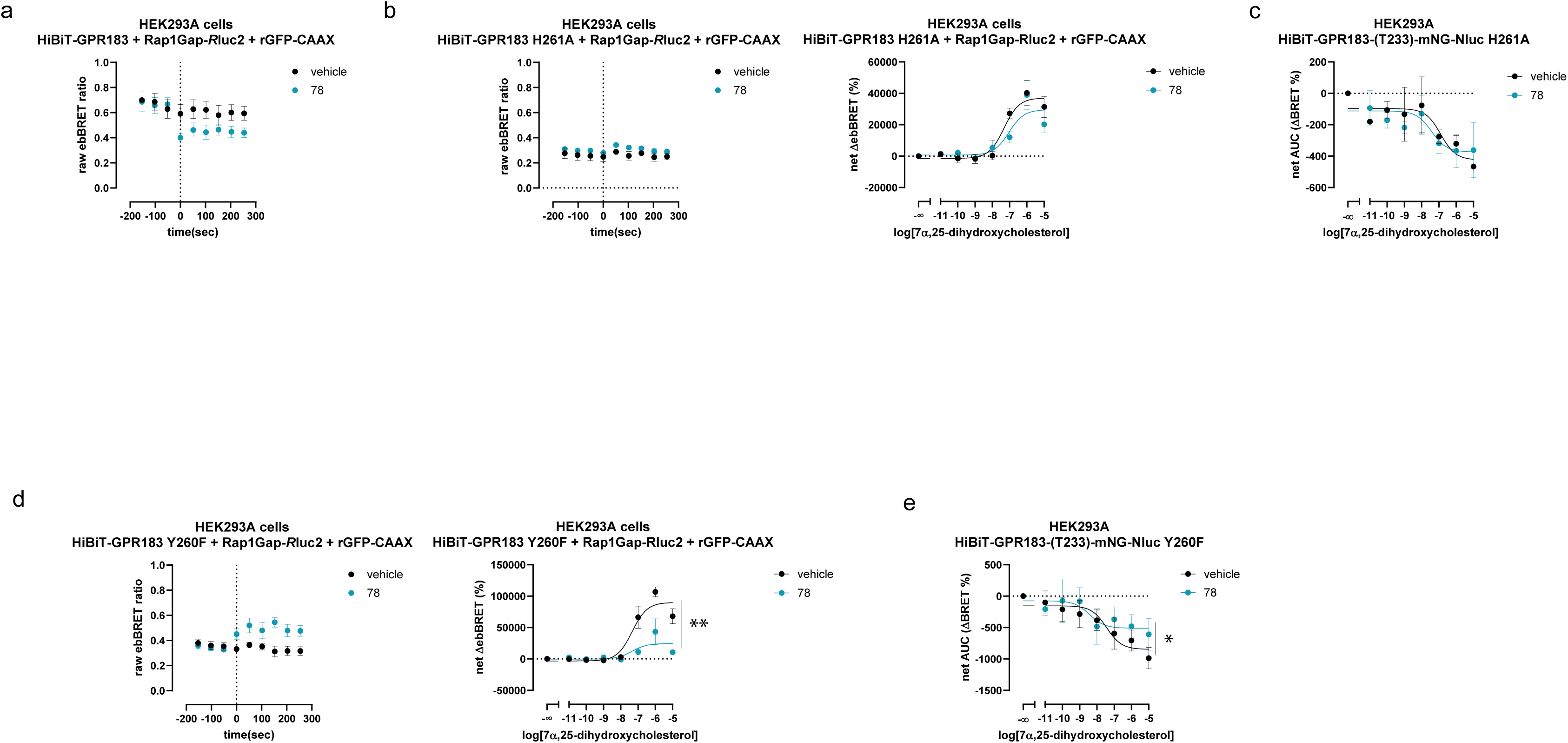
The key interactome of 78 in the GPR183: *in vitro* analysis. **a)** Addition of 10 μM of **78** (time = 0, marked as dashed lines) leads to a reduction in the ebBRET signal indicative of a decrease in GPR183-mediated Gi activation. The data are presented as raw ebBRET ratio; vehicle (1% DMSO) was not subtracted. **b) Left**: Addition of 10 μM of **78** did not modify constitutive GPR183 H261A^6.52^-mediated Gi activation. The data are presented as raw ebBRET ratio; vehicle (1% DMSO) was not subtracted. **Right**: Preincubation with 10 μM of **78** did not modify agonist-simulated GPR183 H261A^6.52^-mediated Gi activation. The data are presented as net AUC of ΔebBRET % (over the three baseline measurements); vehicle (1% DMSO) was subtracted). **c)** The concentration response curve of 7α,25-dihydroxycholesterol on the HiBiT-GPR183-(T233)-mNG-Nluc H261A^6.52^ in the presence or absence (vehicle, 1% DMSO) of 10 μM of **78**. **78** or vehicle were added, and the BRET signal was measured over 10 minutes, followed by the addition of different concentrations of 7α,25-dihydroxycholesterol. The plate was measured for another 15 minutes, but since the mutations also impact the agonist binding kinetics, only the first three minutes were used in the analysis. The data are presented as net AUC of ΔBRET % (over the 10-minute-long read); vehicle (1% DMSO) was subtracted. The differences in agonist potency and BRET_max_ were assessed by comparing log*EC*_50_ and top values, respectively, for the two curves with the extra sum-of-squares F test. **d) Left**: Addition of 10 μM of **78** induced GPR183 Y260F^6.51^-mediated Gi activation. The data are presented as raw ebBRET ratio; vehicle (1% DMSO) was not subtracted. **Right**: Preincubation with 10 μM of **78** did impaired agonist-simulated GPR183 H261A^6.52^-mediated Gi activation. The data are presented as net AUC of ΔebBRET % (over the 10-minute-long read); vehicle (1% DMSO) was subtracted). The differences in ebBRET_max_ were assessed by comparing top values for the two curves with the extra sum-of-squares F test, and the difference is indicated with the line and marked for statistical significance. **e)** The concentration response curve of 7α,25-dihydroxycholesterol on the HiBiT-GPR183-(T233)-mNG-Nluc Y260F^6.51^ in the presence or absence (vehicle, 1% DMSO) of 10 μM of **78**. **78** or vehicle were added, and the BRET signal was measured over 10 minutes, followed by the addition of different concentrations of 7α,25-dihydroxycholesterol. The plate was measured for another 15 minutes, but since the mutations also impact the agonist binding kinetics, only the first three minutes were used in the analysis. The data are presented as net AUC of ΔBRET % (over the first 10-minute-long read); vehicle was subtracted. The differences in agonist potency and BRET_max_ were assessed by comparing log*EC*_50_ and top values, respectively, for the two curves with the extra sum-of-squares F test, and the difference is indicated with the line and marked for statistical significance. The data in this figure are shown as mean ± s.e.m. of three to four independent experiments.

Interestingly, in the Y260F^6.51^ mutant, the addition of 10 µM **78** increased the basal GPR183-mediated Gi activation (**Figure 7d**). In line with this, co-application of various concentrations of 7α,25-dihydroxycholesterol in the presence of **78** resulted in reduced potentiation of Gi activation compared to vehicle control (ebBRET_max_, P = 0.002), while the agonist’s potency remained unchanged (p*EC*₅₀ = 7.10 (95% CI: 6.21–8.40) vs. p*EC*₅₀ = 7.37 (95% CI: 7.02–7.73); P = 0.62; **Figure 7d**). Similarly, **78** did not significantly alter the potency of the agonist to induce the active conformation of GPR183 (p*K*_d_ = 8.43 (95% CI: 6.44–9.65) vs. p*K*_d_ = 7.44 (95% CI: 5.79–9.23) for **78** and vehicle, respectively; P = 0.22), but did reduce BRET_max_ (P = 0.02), suggesting that **78** stabilizes a distinct receptor conformation at saturating agonist concentrations (**Figure 7e**). Together, these data indicate that **78** may in fact function as a partial agonist in the GPR183 Y260F^6.51^, potentially stabilizing a different conformation of the receptor (**Figure S5**), where it competes with the orthosteric agonist and thereby reduces its maximal efficacy due to a ceiling effect.

Collectively, these findings validate the accuracy of our MD-based predictions and underscore the critical roles of H261^6.52^ and Y260^6.51^ in anchoring **78** within the GPR183 orthosteric binding pocket. It has to be noted that, consistently with the literature [11, 54], these two mutations also impaired the pharmacological properties of the agonist.

### Compound 78 inhibits GPR183-mediated lymphocyte migration, particularly of CD4+T and B cells

We investigated the effect of compound **78** on primary lymphocyte migration using an in *vitro* migration assay (**Figure 8a**). Stimulation with 7α,25-dihydroxycholesterol induced lymphocyte migration (P<0.05), with a magnitude similar to that observed with the positive control CCL5 (P<0.05), in comparison to unstimulated conditions (US) (**Figure 8b**). Next, we performed a sub-analysis of the transmigrated cell compartments. Stimulation with 7α,25-dihydroxycholesterol induced transmigration of CD4+ T cells and B cells (P<0.05), while the addition of CCL5 only promoted migration of T (CD4+ and CD8+) cells (P<0.05) (**Figure 8c**). This is consistent with the expression of GPR183 on CD4+ and B cells (**Figure S6a**).

**Figure 8.**
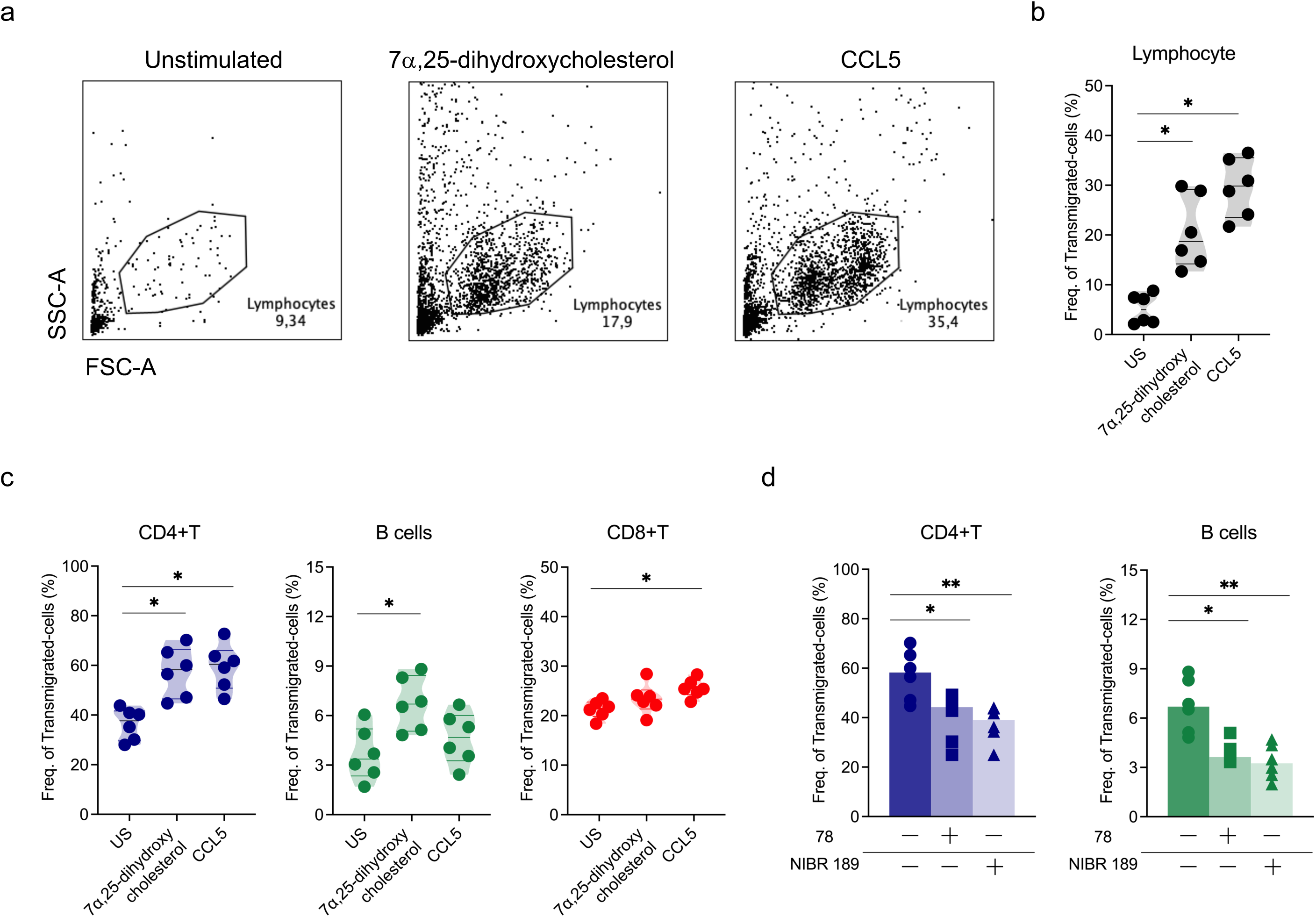
Compound 78 inhibits GPR183-mediated lymphocyte, particularly CD4+T and B cells, migration. **a)** Flow cytometric plots representing transmigrated lymphocytes of one healthy control (HC) after stimulation with 7α,25-dihydroxycholesterol (0.1% DMSO final concentration) and CCL5, and unstimulation (US). Frequencies of migratory **b)** lymphocytes and **(c)** CD4+T, B and CD8+T cells in HC (n=6) after stimulation. **d)** Frequencies of transmigrated CD4+T (left panel) and B (right panel) cells after treatment with compound **78** and NIBR189 in 7α,25-dihydroxycholesterol-stimulated samples from HC (n=6). All comparisons were analysed by Kruskal-Wallis test and P values were corrected by Dunn’s test for multiple comparisons. Only statistically significant P values < 0.05 are presented.

The addition of compound **78** reduced the proportion of transmigrated lymphocytes to the same level as NIBR189 (**Figure S6b, left panel**), as compared to the conditions without inhibitor (P<0.05). Of note, treatment with these compounds did not significantly affect CCL5-mediated migration **(Figure S6b)** nor cell viability **(Figure S6c)**. Furthermore, this equivalent effect was observed across specific cell types as both compound **78** and NIBR189 reduced the migration of CD4+T and B cells but had no impact on CD8+T cells (**Figure 8d and Figure S6d).** These data conclusively show that compound 78 inhibits GPR183-mediated primary lymphocyte, particularly CD4+T and B cells, migration to the same high degree of efficacy as NIBR189.

## DISCUSSION

In this study, we report the discovery of a first-in-class GPR183 inverse agonist, identified through an integrated AI-driven SBVS strategy and validated using several biophysical tools, including an innovative conformational BRET-based biosensor. GPR183, a Gi-coupled chemotactic receptor with roles in immune cell trafficking and disease pathogenesis, remains an underexploited drug target despite mounting evidence linking its activity to various inflammatory, autoimmune, and malignant conditions. Previous efforts to pharmacologically target GPR183 have focused on piperazine diamides and benzo[d]thiazole-based inverse agonists. Our study aimed at expanding the chemical repertoire for GPR183 ligands and identified a series of pyrazolpyridazinones, among which the most interesting one, compound **78**, functions as a GPR183 inverse agonist with balanced activity on both Gi and β-arrestin pathways.

Here, the discovery pipeline leveraged PyRMD2Dock, a hybrid AI-docking platform designed to enrich VS hits with favourable predicted binding profiles and contact maps aligned with key GPR183 residues. This approach proved effective in identifying promising binders from a large chemical space, yielding a >20% initial hit rate from the primary 70-compound screen, and subsequently leading to the selection of compound **43** and its more efficacious analogue, compound **78**. Functional assays revealed that compound **78** significantly reduces both constitutive and agonist-induced Gi signaling and impairs agonist-stimulated β-arrestin2 recruitment to overexpressed GPR183. MD simulations, substantiated by mutagenesis data, further validated the predicted binding mode, highlighting key interactions with residues e.g. Y260^6.51^ and H261^6.52^, which appear essential for ligand engagement and receptor stabilisation in an inactive conformation. To this end, it is the first study that showed a successful application of PyRMD2Dock for GPCRs.

Another technological advancement of this study is the development of a conformational biosensor for GPR183, constructed by fusing mNG into the ICL3 and Nluc at the C-terminus of the receptor. Among several sensor configurations tested, the T233 insertion site yielded visibly the most robust and reliable BRET signal changes upon agonist stimulation. This tool enabled us to directly monitor real-time, ligand-induced conformational rearrangements of GPR183 in live cells. Notably, the sensor responded to the endogenous agonist 7α,25-dihydroxycholesterol in a concentration-dependent manner, permitting estimation of the binding affinity. This pharmacological parameter was in good agreement with previous ligand binding studies, underscoring the utility of this biosensor for mechanistic studies (literature: 7α,25-dihydroxycholesterol p*K*_d_ = 9.34 [8]; our study: p*K*_d_ = 9.15). Next, the sensor allowed the competition binding studies and analysis of NIBR189 and **78** binding. We have already discussed that the conformational sensors have not been really explored for orphan GPCRs [55], with only a very recent pre-print introducing a Halo-Nluc-based sensor of GPR3 (Schihada H, et al., doi:10.26434/chemrxiv-2025) [56]. To this end, a GPR183-oriented sensor based on a similar Halo-Nluc principle was not functionally responding to an agonist simulation, which underlines the need for a careful tool design for GPCRs.

While potent inverse agonists such as NIBR189 represent significant advances, their shared piperazine diamide scaffold might suffer from inherent liabilities, including metabolic instability. Such highly optimized scaffolds often reach a developmental plateau, where improving one property (e.g., metabolic stability) negatively impacts another (e.g., potency or selectivity). In this context, the discovery of **78** might unlock a new opportunity for drug design, unencumbered by the constraints of the previous scaffold. Interestingly, the compound seems to act as a partial agonist when the binding site is modified to include F260^6.51^ in the place of Y260^6.51^. These findings will likely contribute to deepening our understanding of the activation mechanism of GPR183. Likewise, structural novelty of **78** provides the foundation for developing new GPR183 modulators – in principle inverse agonists but, given the results of the GPR183 Y260F^6.51^ assays, also agonists – possibly even enabling the engineering of additional biased ligands for GPR183 [57], the optimization of drug-like properties, and the establishment of a robust safety profile from a superior starting point. Importantly, the capacity of compound **78** to inhibit GPR183-dependent cell migration in lymphocytes, with notable effects on CD4+ T and B cells, underscores its potential functional significance in immune-related contexts and provides a rationale for its therapeutic exploration in inflammatory and autoimmune diseases.

In summary, this work combines computational innovation, chemical biology, and receptor pharmacology to deliver a novel GPR183 modulator and a new biosensor system. Together, they provide the foundation for a more nuanced understanding of GPR183 signaling and pharmacology, and open new avenues for the discovery of therapeutic agents targeting this and related GPCRs in immune and inflammatory diseases.

## STUDY LIMITATIONS

Nevertheless, several limitations should be acknowledged.

First, the efficacy, potency and affinity of compound **78**, while sufficient for proof-of-concept studies, represent a promising starting point rather than a fully optimized profile when compared to benchmark compounds such as NIBR189. The half-maximal inhibitory concentration values of **78** in both Gi and conformational sensor assays (re-calculated to *K*_i_) remain in the high nanomolar/ low micromolar range, which highlights the need for further chemical optimisation to unlock its potential for *in vivo* applications or therapeutic translation.

Next, even though this study does not focus specifically on the agonist-stimulated GPR183-mediated responses, but rather on the inverse agonist discovery, we have concluded that the following has to be appreciated: literature reports advise that a 1% final DMSO concentration be present with working dilutions of the 7α,25-dihydroxycholesterol [9]. Although we did not observe visible precipitation of the agonist at its highest concentrations with 0.1% DMSO in our assays, we used 1% DMSO in the majority of experiments to ensure solubility and reproducibility. Still, this solvent concentration appears to alter BRET baselines in the conformational sensor readout, leading to elevated variability in the vehicle controls and consequently non-zero standard deviations. This variability negatively impacts Z-factor calculations for the experiments where 1% DMSO was employed (not shown here explicitly but can be deducted from e.g. **Figure 4b**). To this end, for GPR183, the mNG-Nluc sensor demonstrates responsiveness to ligands, it allows important mechanistic insights into the receptor and shows a very significant improvement over a Halo-Nluc-based GPR183 sensor. However, it does not *per se* currently meet the quantitative robustness criteria needed for high-throughput screening (Z-factor > 0.5). On the other hand, the G protein-coupling ability of our sensor is likely compromised which, together with a currently-understood ligand-stimulated activation mechanism of Gi-coupled receptors with the TM6 swing-out ca. 3.0Å smaller than e.g. for Gs-coupled receptors [58], could presumably lead to and explain more moderate ΔBRET .

## FINAL CONSIDERATIONS

Looking ahead, this study provides a clear foundation for several exciting opportunities. The validated computational models and structure-activity relationships established for compound **78** now offer a rational platform for designing next-generation derivatives. Further exploration, directly informed by our docking and MD predictions, will yield compounds with significantly enhanced potency and optimized pharmacological properties.

Simultaneously, our work has identified clear strategies to improve the conformational sensor’s performance for high-throughput applications. Focused efforts on refining the assay - through linker optimization, fluorophore variants, and assay miniaturization - can unlock its full potential for high-content or high-throughput applications. Furthermore, our findings highlight that the design of more soluble GPR183 agonists would be a valuable contribution to the entire field.

Beyond its immediate application for GPR183, the conformational sensor described here provides a versatile and readily adaptable platform. This design should be applicable to the studies of other GPCRs, many of which are high value but challenging targets that remain poorly understood and lack selective synthetic ligands. Thus, our work delivers not only specific insights into GPR183 but also a powerful tool to accelerate discovery across a landscape of difficult yet relevant drug targets.

## DATA AVAILABILITY

All data sets generated and analysed during the current study are available from the corresponding author on reasonable request.

## AUTHOR CONTRIBUTIONS

Andersson L was responsible for methodology, investigation, data curation, and writing. Roggia M contributed to writing, methodology, investigation, and data curation. Wangriatisak K contributed to writing, investigation, and data curation. Gil M contributed to investigation. Chemin K was responsible for writing, supervision, and funding acquisition. Cosconati S was responsible for conceptualization, methodology, writing, supervision, and funding acquisition. Kozielewicz P was responsible for conceptualization, investigation, writing, supervision, project administration, methodology, and data curation.

## ASSOCIATED CONTENT

### Supporting Information

**Table S1.** Antibodies used for chemotaxis assay.

| Antibody | Fluorophore | Clone | Company | Dilution | Catalogue No. |
| --- | --- | --- | --- | --- | --- |
| Fixable viability dye | Green | - | Biolegend | 1:2000 | 423107 |
| CD3 | PE/Cy5.5 | 7D6 | ThermoFisher | 1:100 | MHCD0318 |
| CD4 | BV786 | SK3 | BD | 1:100 | 563877 |
| CD8 | BUV395 | RPA-T8 | BD | 1:100 | 563795 |
| CD14 | FITC | M5E2 | BD | 1:100 | 555397 |
| CD16 | APC/H7 | 3G8 | BD | 1:200 | 560195 |
| CD19 | BV421 | HIB19 | Biolegend | 1:50 | 302234 |
| CD56 | PE/Cy5 | B159 | BD | 1:100 | 555517 |
| GPR183 | APC | SA313E4 | Biolegend | 1:50 | 368908 |

**Figure S1.**
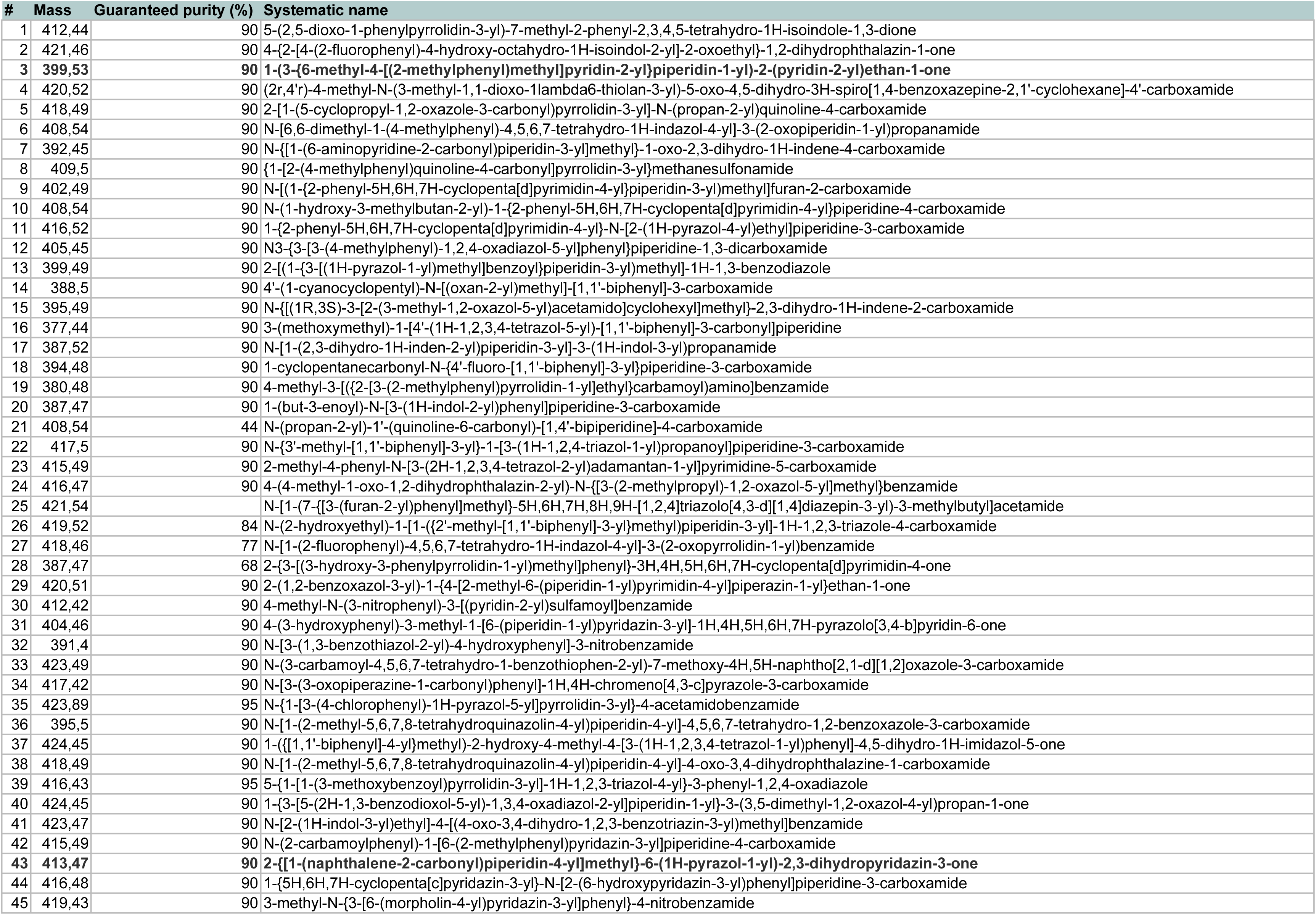
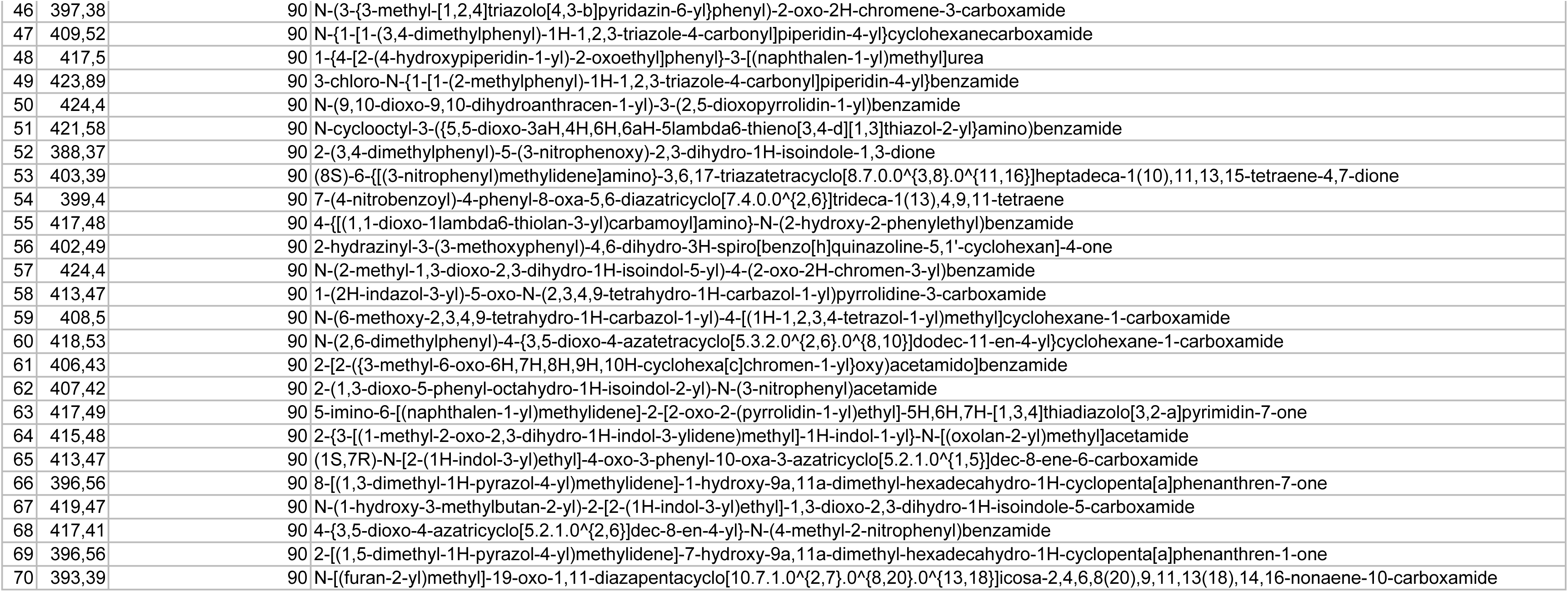
List of the compounds selected from the VS.

**Figure S2.**
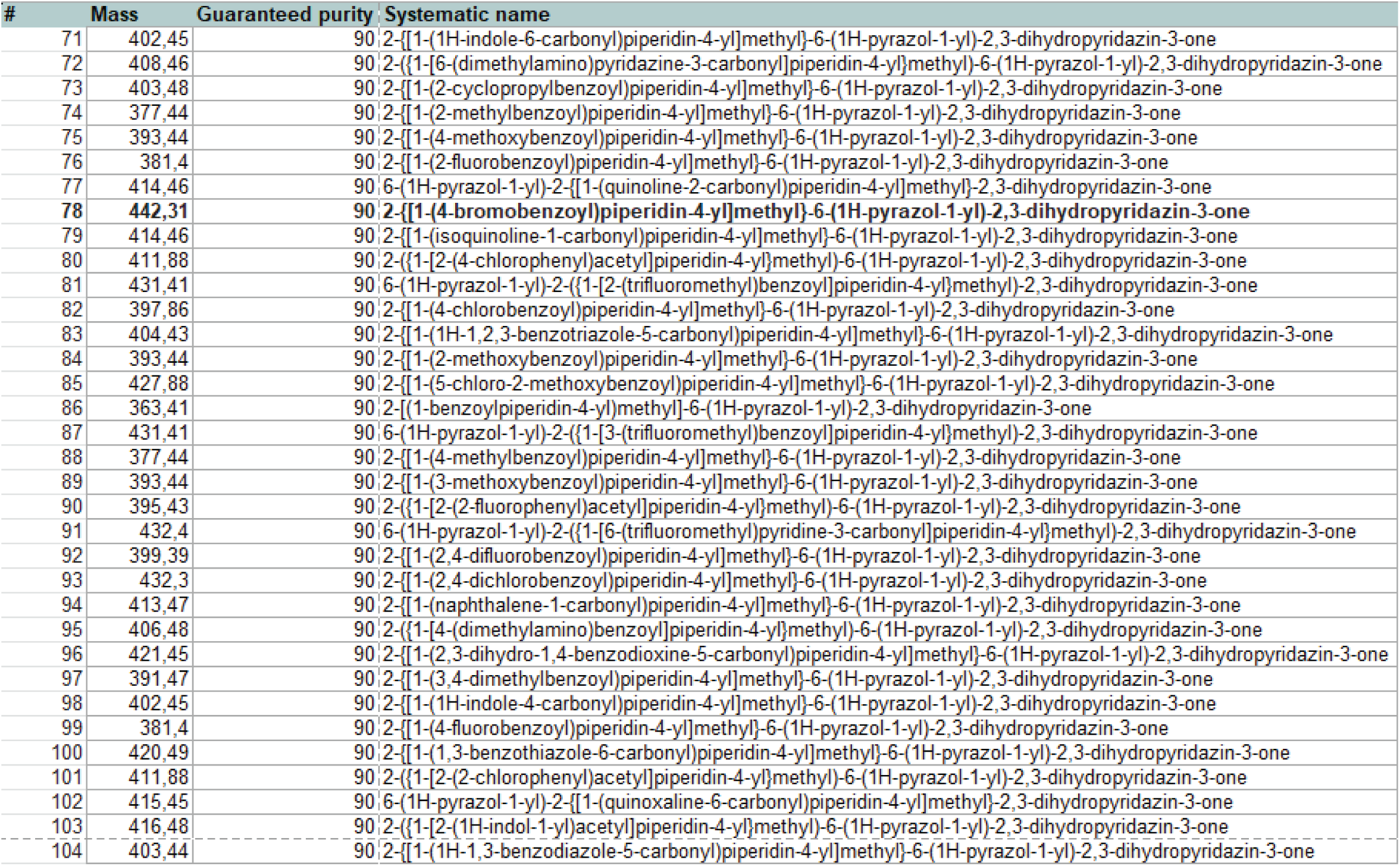
List of the compound selected from the hit expansion.

**Figure S3.**
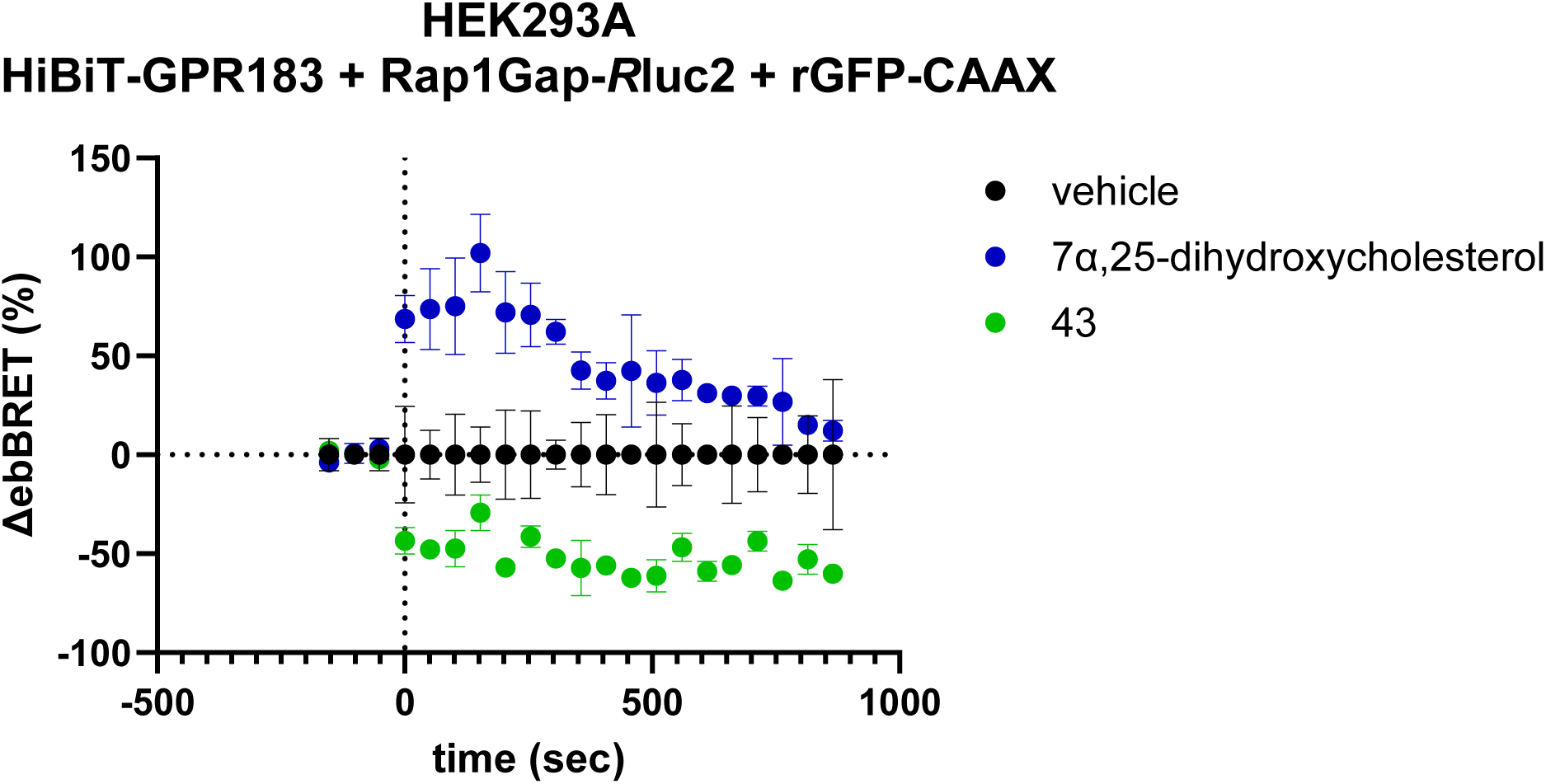
The kinetic plot from a representative ebBRET-based Gi activation assay. The plate was measured three times to obtain baseline BRET, and then the compounds or the vehicle (0.1% DMSO) were added, and the plate was measured for another 15 minutes. The data are presented as ΔebBRET % over the three baseline reads measured prior to the compound/vehicle addition; vehicle (0.1% DMSO) was subtracted. The data are shown as mean ± s.d. of two technical replicates. The data were analysed for differences using Mann-Whitney test; P = 0.4.

**Figure S4.**
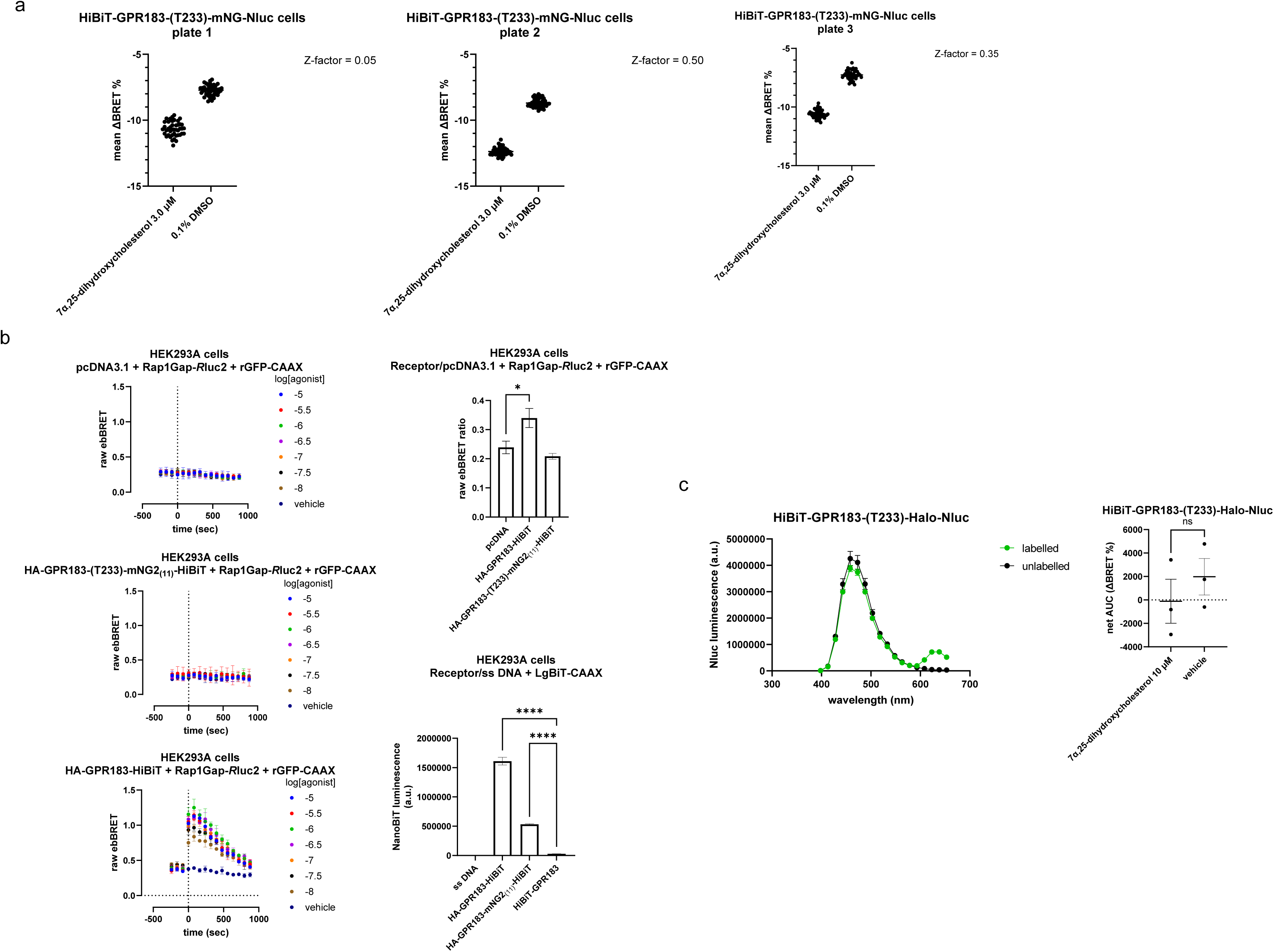
**a)** Z-factor evaluation of the HiBiT-GPR183-(T233)-mNG-Nluc sensor using the HEK293A cells stably overexpressing the construct. The data are presented as the mean ΔBRET% (over the three initial baseline reads) from the 15-minute-long read following the addition of the agonist or vehicle (0.1% DMSO) for three separate plates. **b) Left top**: pcDNA.3.1-transfected cell nor (**left middle**) HA-GPR183-(T233)-mNG2_(11)_-transfected cells do not respond to agonist stimulation to activate Gi in GPR183-specific manner as opposed to (**left bottom**) HA-GPR183-HiBiT, which does not have an ICL3-inserted tag. 0.1% DMSO was used as a vehicle. The data come from a representative experiment of three independent experiments and are shown as mean ± s.d. of three technical replicates. **Right top**: Ligand-unbound HA-GPR183-(T233)-mNG2_(11)_ does not constitutively activate Gi. **Right bottom**: HA-GPR183-(T233)-mNG2_(11)_ is trafficked to the cell membrane. The data are shown as mean ± s.e.m. of three independent experiments. The data were analysed for differences using one-way ANOVA with multiple comparison Dunnett’s post-hoc analysis; * P<0.05, ****P < 0.0001. **c) Left**: Upon the overexpression of the HiBiT-GPR183-(T233)-Halo-Nluc conformational sensor and addition of the Halo substrate, there is an energy transfer between the Nluc and Halo (detectable a luminescence peak over 600 nm). **Right**: The addition of 10 μM of the agonist does not result in a notable change in the BRET signal in comparison with the vehicle (0.1% DMSO).

**Figure S5.**
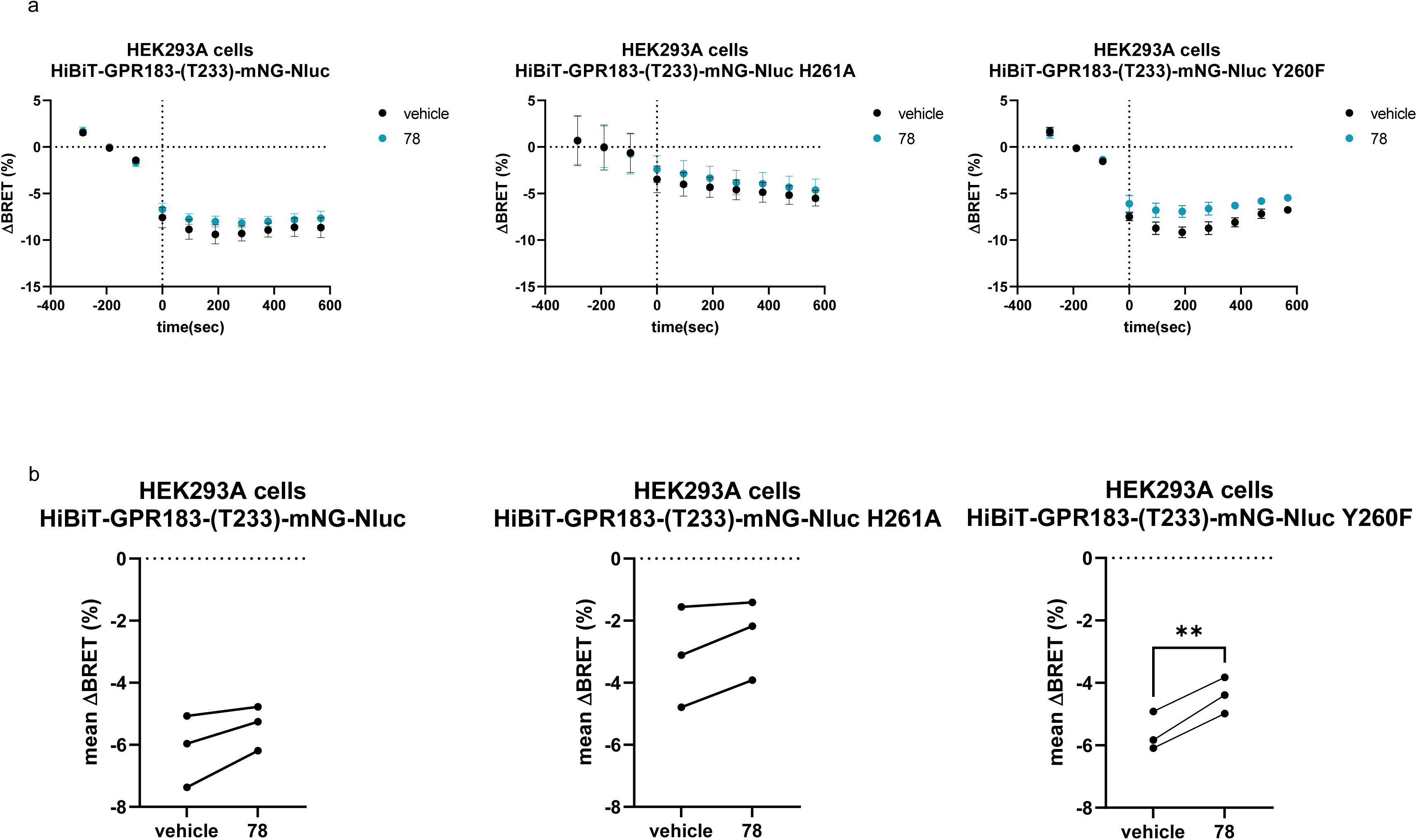
**a)** Addition of 10 μM of **78** leads to a subtle but visible change in BRET in HiBiT-GPR183-(T233)-mNG-Nluc Y260F^6.51^ conformational biosensor in comparison to vehicle (1% DMSO). This is further supported by the statistical analysis in **(b)**. The data are presented as the mean ΔBRET% (over the three initial baseline reads) from the 10-minute-long read following the addition of **78** or vehicle. The data are shown as mean ± s.e.m. of three independent experiments. The data in **b)** were analysed with a paired *t*-test; ** P<0.01

**Figure S6.**
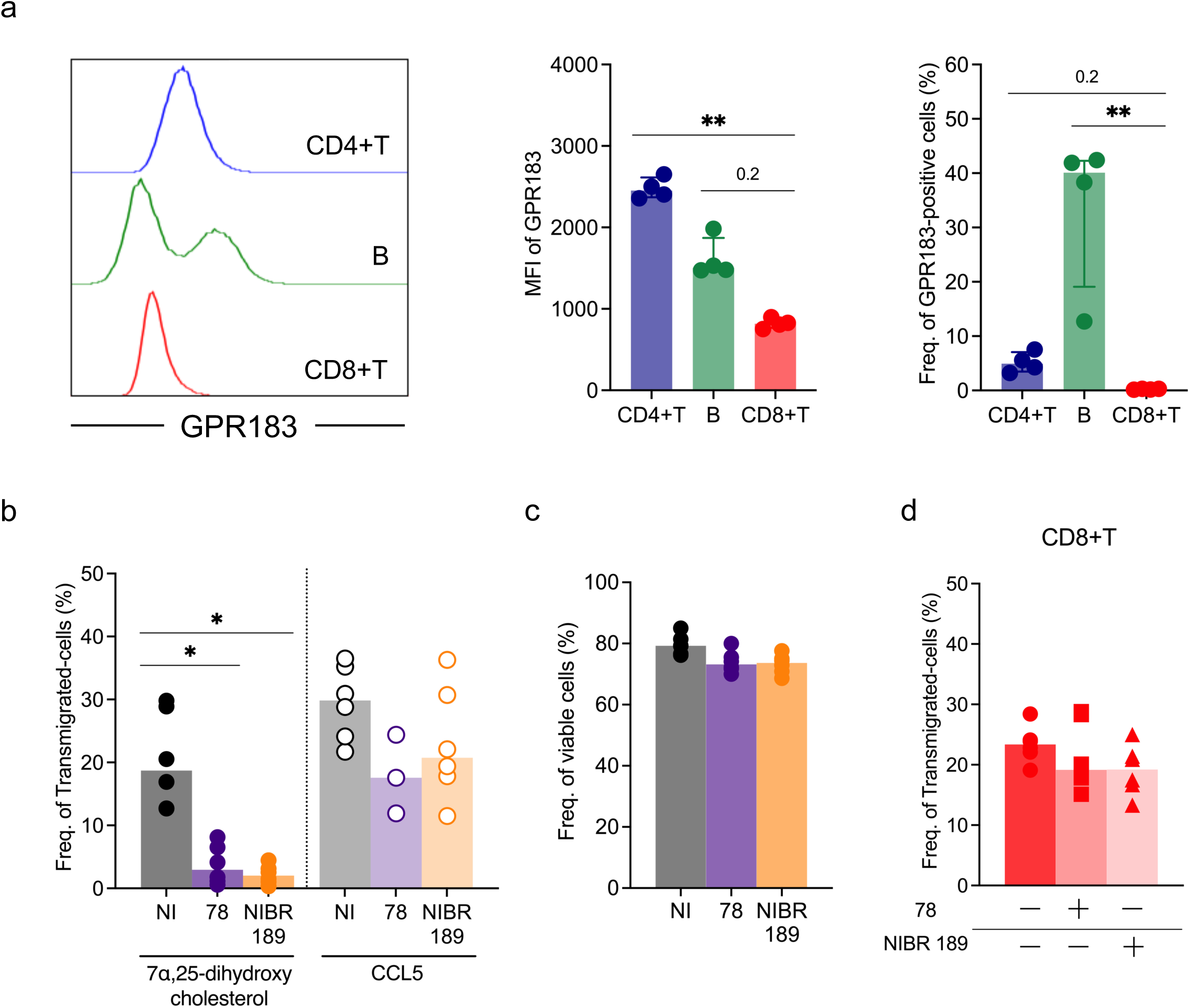
Histogram overlays representing GPR183 expression. (**a, left panel**), median fluorescent intensity (MFI) of GPR183 (middle panel), and frequency of GPR183-positive cells (right panel) of GPR183 in CD4+T, B and CD8+T cells in healthy control (HC, n=4). **b)** Frequency of migratory lymphocytes treated with **78**, NIBR189, or no-inhibitor (NI) after stimulation in HC. **c)** Frequency of viable cells treated with **78**, NIBR189, or NI following 7α,25-dihydroxycholesterol stimulation. **d)** Frequency of transmigrated CD8+T cells after addition of **78** and NIBR189 to 7α,25-dihydroxycholesterol-stimulated (0.1% DMSO final concentration) samples from HC.

## ACKNOWLEDGMENTS

Kozielewicz P thanks Prof. Gunnar Schulte for his support.

## ABBREVIATIONS

GPCR: G protein-coupled receptor
AI: Artificial Intelligence
MD: molecular dynamics
VS: virtual screening.

